# The PBAF chromatin remodeling complex contributes to metal homeostasis through Mtf1 regulation

**DOI:** 10.1101/2025.04.12.648552

**Authors:** Nick Carulli, Emma E. Johnston, David C. Klein, Odette Verdejo-Torres, Anand Parikh, Antonio Rivera, Michael Quinteros, Aidan T. Pezacki, Christopher J. Chang, Sarah J. Hainer, Teresita Padilla-Benavides

## Abstract

SWI/SNF chromatin remodeling complexes regulate gene expression by modulating nucleosome positioning, yet their roles in metal homeostasis during skeletal muscle development remain unclear. Here, we uncover distinct functions of the BAF, PBAF, and ncBAF complexes in myoblast proliferation under metal stress. While knockdown (KD) of *Baf250a* (BAF-specific) or *Brd9* (ncBAF-specific) reduces myoblast proliferation, *Baf180* (PBAF-specific) KD does not impair cell proliferation under basal conditions. Interestingly, supplementation with copper (Cu) or zinc (Zn) rescues proliferation in *Baf250a*- and *Brd9*-deficient myoblasts but paradoxically inhibits growth in *Baf180* KD cells. Mechanistically, *Baf180* KD disrupts Cu and Zn homeostasis, leading to intracellular Cu accumulation without labile Cu⁺ pools and impaired expression of *Atp7a*, a key Cu exporter. Transcriptomic analyses reveal widespread gene dysregulation in metal-treated *Baf180*-deficient cells, while metal supplementation promotes pro-proliferative gene expression in *Baf250a*- and *Brd9*-KD myoblasts. CUT&RUN assays demonstrate that metal-responsive transcription factor Mtf1 exhibits increased chromatin binding upon Cu treatment, targeting genes involved in stress response and myogenesis. Notably, Mtf1 colocalizes with Baf180 in the nucleus and co-immunoprecipitates with both conserved SWI/SNF subunits and Baf180, suggesting a functional interplay between PBAF and Mtf1 in regulating metal-dependent gene expression. Our findings establish the PBAF complex as a crucial regulator of Cu/Zn homeostasis in myoblast proliferation via Mtf1, while metal supplementation compensates for BAF and ncBAF dysfunction but exacerbates defects in PBAF-deficient cells. This study reveals a novel link between chromatin remodeling, metal signaling, and muscle development, with implications for stress adaptation and metabolic regulation in myogenesis.

## INTRODUCTION

Mammalian DNA is organized into chromatin, where nucleosomes compact approximately 150 bp of DNA around histone proteins, reducing its accessibility to transcription factors (1, 2). Post-translational histone modifications alter chromatin accessibility, recruiting specific factors and influencing chromatin structure (3–5). ATP-dependent chromatin remodelers, such as SWI/SNF complexes, utilize ATP hydrolysis to reposition or evict nucleosomes, facilitating euchromatin formation (6, 7). All mammalian SWI/SNF complexes contain an ATPase subunit, either Brahma (BRM) or Brahma-related gene 1 (Brg1) (8–10). These complexes include the canonical cBAF (BRG1/BRM-associated factor), PBAF (polybromo-associated BAF), and ncBAF (non-canonical BAF) complexes, each with distinct subunit compositions (11). The BAF complex contains Baf250a and Dpf2, PBAF includes Baf180, Baf200, and Brd7, while ncBAF is characterized by Brd9 and GLTSCR1 (11–14). These complexes exhibit specialized roles depending on developmental stage and cellular lineage, many of which remain incompletely understood. For instance, in glucocorticoid receptor (GR)-mediated chromatin remodeling in SW-13 cells, Brg1 and Baf250a, but not Baf180, were required (15, 16). In this sense, Brg1 was shown to play a critical role in supporting the transcriptional response to GR activation, and its disruption significantly shifts the landscape of hormone-induced gene expression. Notably, GR binding sites that are pre-occupied by Brg1 show an enrichment for motifs recognized by pioneer factors such as FOXA1 and GATA3 (17). BAF60c is a chromatin remodeling component involved in insulin-induced lipogenic gene transcription in the liver (18). In mice, Baf60c plays a key role in early embryonic development, particularly in heart and somite formation (19). RNA interference (RNAi)-mediated Baf60c silencing disrupts heart morphogenesis, anterior/secondary heart field expansion, and cardiac and skeletal muscle differentiation (19). Partial Baf60c reduction mimics congenital heart defects, whereas overexpression facilitates interactions between cardiac transcription factors and Brg1, enhancing target gene activation (19).

Similarly, Baf250a is essential for cardiac progenitor development, as its ablation in the second heart field causes embryonic lethality and cardiac defects. In embryonic stem cells, *Baf250a* deficiency impairs cardiomyocyte formation by disrupting key regulators such as *Mef2c*, *Nkx2.5*, and *Bmp10* (20). Brd9, a defining ncBAF subunit, interacts with androgen receptors (AR) to modulate AR-dependent gene expression in prostate cancer cells, where it is required for survival (21). Chromatin mapping studies in synovial sarcoma and malignant rhabdoid tumors indicate that ncBAF uniquely localizes to CTCF sites and promoters. Brd9 depletion impairs proliferation in these cancer cells (22). Conversely, knockout (KO) of *Baf180*, a PBAF subunit, in mouse embryonic fibroblasts leads to premature senescence and disrupted hematopoiesis (23). Baf180 regulates the expression of *p21*, a cyclin-dependent kinase inhibitor, by binding its promoter and repressing transcription, thereby controlling cell cycle progression (24). Actively transcribed genomic regions are prone to genomic instability, and transcription is often suppressed near DNA double-strand breaks (DSBs) (25). The PBAF complex silences transcription at DSB flanking regions and aids in repair, a process dependent on ataxia-telangiectasia mutated (*ATM*) kinase and Baf180 phosphorylation (25). PBAF also enhances transcriptional activation by nuclear receptors, including retinoic acid receptor-α (*RXRα*), vitamin D receptor (*VDR*), and peroxisome proliferator-activated receptor γ (*PPARγ*) (26). Baf180 deletion in mouse embryos results in severe ventricular hypoplasia and trophoblast placental defects, mirroring RXRα and PPARγ deficiencies, with heart defects directly attributable to *Baf180* loss (26). Target gene analysis suggests that BAF180 regulates cardiac chamber maturation by modulating genes such as *S100A13*, *RARβ2*, and *CRABPII* (26).

Our group has previously reported on the roles of BAF, ncBAF, and PBAF complexes in skeletal muscle lineage (12, 13, 27). Using subunit-specific (*Baf250a*, *Brd9*, and *Baf180*) shRNA-mediated knockdown (KD) in C2C12 myoblasts, we reported that the BAF complex is essential for myoblast proliferation and differentiation by regulating the expression of *Pax7* and *Myogenin* to sustain the myogenic program (12, 13). Mechanistic studies revealed that Baf250a binds regulatory sequences of these genes during proliferation and differentiation. While ncBAF influences RNA metabolism and indirectly contributes to myogenesis, PBAF appears dispensable for these processes (12, 13, 28). However, *Baf180* KD resulted in impaired expression of stress-related genes, particularly those involved in metal transport and homeostasis (12).

Metal ions such as copper (Cu) and zinc (Zn) play crucial roles in myogenesis by supporting mitochondrial respiratory chain complexes and enzymatic activities. Cu is essential for myoblast proliferation and differentiation (29, 30). Mtf1, a Zn-finger transcription factor responsive to metals including Cu, is required for myoblast differentiation (31). PiC2, a Cu transporter regulated by Mtf1, is upregulated during myogenesis and is necessary for proper proliferation and differentiation (31, 32), and Crip2, a novel Cu-binding transcriptional regulator, is also required for myogenesis and maintenance of metal homeostasis (33).

Both SWI/SNF complexes and metal regulatory networks are integral to myogenesis (12, 13, 28, 30–35). In this study, we explored the contributions of BAF, PBAF, and ncBAF complexes to metal regulation and homeostasis using shRNA-mediated KD in C2C12 cells. Our data demonstrates that Zn or Cu supplementation rescues proliferation defects in C2C12 myoblasts with dysfunctional BAF or ncBAF complexes. Additionally, PBAF is required for metal regulation in proliferating myoblasts, as *Baf180* KD cells exhibit impaired proliferation upon Zn and Cu supplementation. These findings highlight an emerging role of SWI/SNF complexes in metal homeostasis during myogenesis.

## MATERIALS AND METHODS

### Antibodies

Hybridoma supernatant against Pax7, obtained from the Developmental Studies Hybridoma Bank (University of Iowa; deposited by A. Kawakami), was used for immunocytochemistry. Normal rabbit IgG (sc-2027) and mouse anti-Mtf1 (sc-365090) were obtained from Santa Cruz Biotechnologies. Rabbit anti-Baf180 (A0334), rabbit anti-Baf250a (A16648) and anti-GAPDH (A19056) were from Abclonal. Rabbit anti-Brd9 (PA5-113488) was from Thermo Fisher Scientific. Rabbit anti-Baf180 (89123S), -Baf250a (12354S), and -Brd9 (48306S), used for confocal microscopy at dilutions of 1:50, 1:50, and 1:100 respectively, were from Cell Signaling Technology. The secondary antibodies were HRP-conjugated anti-rabbit IgG (31460), goat anti-rabbit IgG Alexa Fluor Plus 594, goat anti-mouse IgG Alexa Fluor Plus 488, goat anti-rabbit IgG Alexa Fluor Plus 633, donkey anti-mouse Alexa-Fluor 594 (A32740, A32723, A21070 and A32744, respectively) were obtained from Thermo Fisher Scientific.

### C2C12 myoblasts culture

Immortalized murine myoblast C2C12 and HEK293T cells were purchased from American Type Culture Collection (ATCC) and cultured in proliferation media composed of Dulbecco’s modified Eagle’s medium (DMEM) supplemented with 10% fetal bovine serum (FBS) and 1% penicillin–streptomycin. Cells were maintained in a humidified incubator at 37°C with 5% CO_2_ at sub-confluent densities.

### Virus production, and transduction of C2C12 myoblasts

Mission plasmids encoding shRNAs targeting specific subunits of mSWI/SNF complexes were obtained from Sigma and used as previously reported (12, 13). *Baf250a* was selected for the BAF complex, and *Baf180* and *Brd9* for the PBAF and ncBAF, respectively (**Supp. Table 1**). The shRNA constructs (15 µg), along with packaging vectors pLP1 (15 µg), pLP2 (6 µg), and pSVGV (3 µg), were transfected into HEK293T cells using lipofectamine 2000 (Thermo Fisher Scientific) following the manufacturer’s instructions. The next day, the culture medium was replaced with 10 mL of fresh DMEM enriched with 10% FBS. The viral supernatant was collected at 24 and 48 h and passed through a 0.22 µm syringe filter (Millipore). For myoblast infection, 5 ml of the filtered supernatant, supplemented with 8 μg/mL polybrene (Sigma), were utilized to transduce 2 × 10^6^ cells, following established protocols (12, 13). Following an overnight incubation, the infected cells underwent selection in growth media containing 2 μg/mL puromycin (Invitrogen). The stable myoblast population was then sustained in growth media supplemented with 1 μg/ml puromycin.

### Western Blot Analyses

C2C12 myoblasts were washed with PBS and solubilized with RIPA buffer (10 mM Tris pH 8, 1% Triton X-100, 0.1% sodium deoxycholate, 0.1% sodium dodecyl sulfate (SDS), 140 mM sodium chloride) and 100 µL of Complete protease inhibitor cocktail (PIC; Thermo Scientific). Samples were sonicated for 12 cycles and protein was quantified via Bradford assay (36). Samples (20 µg) were separated and resolved via SDS-PAGE and electro-transferred to a PVDF membrane (Millipore). Specific proteins were identified with the antibodies described above and peroxidase conjugated secondary antibodies followed by chemiluminescent detection (Tanon, Abclonal Technologies).

### Cell proliferation assays

C2C12 myoblasts were initially seeded at a density of 1 × 10^4^ cells/cm^2^ in the presence of increasing concentrations of CuSO_4_ (50 µM, 100 µM, 200 µM, 300 µM, 500 µM) and ZnSO_4_ (50 µM, 100 µM, 150 µM). Samples were trypsinized at 24, 48, and 72 h post-seeding and cell count, and viability were determined using a Spectrum Cellometer from Nexcelcom Biosciences.

### Immunocytochemistry and light microscopy

C2C12 myoblasts were grown for 48 h in the presence and absence of increasing concentrations of CuSO_4_ (50 µM, 100 µM, 200 µM, 300 µM) and ZnSO_4_ (50 µM, 100 µM, 200 µM). Cells were then fixed overnight at 4°C in 10% formalin in phosphate buffered saline (PBS), washed three times with PBS and permeabilized with Triton X-100 for 10 min. Hybridoma supernatant against Pax7 was used for immunocytochemistry and developed with the universal ABC kit (Vector Laboratories, PK-6200). Images were acquired with an Echo Rebel (Discover Echo) microscope using the 20x objective.

### Immunofluorescence and confocal microscopy

Proliferating C2C12 myoblasts were seeded at 1×10^4^ cells/cm^2^ on coverslips with or without supplementation of 100 µM CuSO_4_ and 50 µM ZnSO_4_ for 48 h. Then, the cells were fixed with 10% formalin in PBS, permeabilized with PBT buffer containing 0.5% Triton X-100 in PBS, and blocked in 5% horse serum in PBT for 1 h. Next, the samples were incubated with the primary antibodies diluted in blocking solution overnight at 4°C. The following day, the myoblasts were washed 3 times with PBT for 10 min at room temperature and incubated with the corresponding secondary fluorescent antibodies in blocking solution for 2 h at room temperature. Nuclei were stained for 30 min with DAPI. Samples were mounted with VectaShield solution (Vector Laboratories) and imaged with a Leica SP8 using the 63X water immersion lens. Images were analyzed with the Leica Application Suite X (Leica Microsystem Inc). Colocalization was quantified via adjusting the Color Threshold of the image and analyzing the particles using Fiji ImageJ software (Version 4.14.0/1.54f) (37).

## CUT&RUN

CUT&RUN was performed as previously described (33, 38–42), using recombinant Protein A-MNase (pA-MNase) (43). Briefly, nuclear extraction was performed on 100,000 cells with a hypotonic buffer (20 mM HEPES-KOH, pH 7.9, 10 mM KCl, 0.5mM spermidine, 0.1% Triton X-100, 20% glycerol and protease inhibitors) and bound to lectin-coated concanavalin A magnetic beads (40 µL beads per 100,000 nuclei; Polysciences). Bead-bound nuclei were chelated with blocking buffer (20 mM HEPES, pH 7.5, 150 mM NaCl, 0.5mM spermidine, 0.1% BSA, 2mM EDTA and protease inhibitors) and washed (wash buffer: 20 mM HEPES, pH 7.5, 150 mM NaCl, 0.5mM spermidine, 0.1% BSA and protease inhibitors). Nuclei were incubated for 1h in wash buffer containing primary antibody (anti-MTF1 (H-6) sc-365090, Santa Cruz Biotechnologies, lot 0050480101; rabbit polyclonal IgG, Abcam ab37415, lot #GR3208186-1), followed by 30-min restriction in pA-MNase diluted in wash buffer, at room temperature with rotation. Samples were equilibrated to 0°C in an ice-water bath and 3 mM CaCl_2_ was added to activate pA-MNase. After 22 min, the digestion reaction was chelated with 20 mM EDTA and 4 mM EGTA, and 1.5 pg MNase-digested *S. cerevisiae* mononucleosomes were added as a spike-in control. Genomic fragments were released after an RNase A treatment and subsequent centrifugation. Isolated fragments were used as input for a library build consisting of end repair and adenylation, ligation of NEBNext stem-loop adapters, and purification with AMPure XP beads (Agencourt). Barcoded fragments were amplified by 15 cycles of high-fidelity PCR and purified using AMPure XP beads. Libraries were pooled and sequenced on an Illumina NextSeq2000 to a depth of ∼10 million uniquely mapped reads.

Paired-end fastq files were trimmed to 25 bp and mapped to the mm10 genome with bowtie2 (options -q -N 1 -X 1000; (44)). Duplicate reads were identified and removed with Picard (45). Reads were filtered for mapping quality (MAPQ ≥ 10) with SAMtools (46) and size classes corresponding to factor-bound footprints (1-120 bp) were generated using a custom awk script (46). Reads were converted to BigWig files using deepTools with RPKM normalization (options - bs 1 --normalizeUsing RPKM; (47)). CUT&RUN peaks were called using HOMER, with IgG-targeted CUT&RUNs used as controls, USING default parameters (48). Heatmaps were plotted using deepTools computeMatrix (options -a 2000 -b 2000 -bs 20 --missingDataAsZero) and plotHeatmap (47). Gene Ontology (GO) analysis was performed on peaks present in at both CUT&RUN replicates using Metascape (49). To control for myoblast-specific background, we kept only genes with a DESeq2 baseMean ≥ 1 (RNA-seq) and used a background list of all genes with a baseMean ≥ 1 in ES cells. P-values were corrected for multiple testing via the Benjamini-Hochberg correction (FDR = 0.05).

### RNA Sequencing (RNA-seq)

RNA-seq was performed as previously described (42). Control scramble (scr) and shRNA transduced myoblasts were treated with 100 µM CuSO_4_ and 50 µM ZnSO_4_, or no metal as detailed above. Cell pellets were then flash-frozen in liquid nitrogen. RNA was extracted from frozen pellets with TRIzol per manufacturer’s instructions and purified by chloroform extraction and isopropanol precipitation. Extracted RNA was flash-frozen in liquid nitrogen and stored until use. Ribosomal RNA was removed from 2 µg of input RNA via antisense tiling oligonucleotides and digestion using thermostable RNase H (MCLabs) (50, 51). rRNA-depleted RNA samples were treated with Turbo DNase (ThermoFisher) and purified by silica column (Zymo RNA Clean & Concentrator kit). rRNA-depleted RNA obtained from each sample was used to build strand-specific RNA-seq libraries using the NEBNext Ultra II Directional Library kit. RNA was fragmented at 94°C for 15 min and used as input for cDNA synthesis and to build strand-specific libraries following the manufacturer’s instructions. Libraries were amplified for 8 cycles of high-fidelity PCR, pooled and sequenced to a depth of approximately 20 million uniquely mapped reads on an Illumina NextSeq2000.

### RNA-seq data analysis

Paired-end fastq files were aligned to the mm10 mouse genome using STAR (options -- outSAMtype BAM SortedByCoordinate --outFilterMismatchNoverReadLmax 0.02 -- outFilterMultimapNmax 1). RNA-seq data was visualized by generating bigwigs files using deepTools with TPM read normalization (options -bs 5 –smoothLength 20 --normalizeUsing BPM)(47). Feature counts were generated using subread featureCounts (options -s 2 -p -B) (52). The count files were imported into R, and subsequent analysis was carried out utilizing DESeq2, implementing the apeglm log_2_ fold change shrinkage correction (53, 54). GO analysis was performed on significantly up- and downregulated genes using Metascape (49). The gene lists utilized for GO analysis comprised exclusively significantly DEG with a DESeq2 baseMean value of ≥ 1, thereby excluding genes with low expression levels. Significance was determined by a DESeq2 adjusted p-value < 0.05.

### Integration of CUT&RUN and RNA-seq datasets

CUT&RUN and RNA-seq datasets were integrated based on gene lists that were called significantly up- or downregulated per DESeq2 results (padj. < 0.05) and CUT&RUN peak files as described above. A bed file containing differentially expressed gene (DEG) promoters was generated using the UCSC Table Browser (55), with promoters defined as regions spanning 1 kb upstream of UCSC RefGene transcription start sites. Direct overlaps between CUT&RUN peaks and DEG promoters were assigned using the HOMER mergePeaks function (options -d given) (48). Overlapping regions were annotated using the HOMER annotatePeaks.pl function and the resulting file was manually separated by specific overlap groups (48). Bed files for each combination of CUT&RUN peaks and DEG promoter regions were generated and used as input for subsequent Gene Ontology, Genome Ontology, and motif enrichment analyses via the HOMER annotatePeaks.pl and findMotifs.pl functions (48). A background gene list of all genes expressed in these experiments (DESeq baseMean value ≥ 1) was used to control for potential overrepresented motif sequences.

### Immunoprecipitation

C2C12 myoblasts were seeded at a density of 1×10^4^ cells/cm^2^ with or without metal solution and incubated for 48 h. Cells were washed three times with PBS, and scrapped off the plates in lysis buffer (50 mM Tris-HCl, pH 7.5, 150 mM NaCl, 1% Nonidet P40, 0.5% sodium deoxicholate, and Complete Protease Inhibitor Cocktail) supplemented with either anti-Mtf1, anti-Baf180 or IgG for 2 h at 4°C, as previously reported (32, 56). PureProteome protein A/G mix magnetic beads (Millipore) were added to each tube and samples were incubated overnight at 4°C on a shaker. The next day, samples were washed three times with 100 mM NaCl in PBS using a magnetic rack to retain the beads. Immunoprecipitated proteins were then eluted from the beads with freshly prepared elution buffer containing 10% glycerol, 50 mM Tris-HCl pH 6.8, and 1 M NaCl. Samples were incubated for 1 h at RT in the elution buffer, and beads were removed using a magnetic rack.

Immunoprecipitated proteins with MTF1 were eluted and subjected to in-solution digestion. Samples were analyzed at the University of Massachusetts Chan Medical School facility. Briefly, samples were reduced with 10 mM dithiothreitol (DTT) at 37°C for 30 min and alkylated with 25 mM iodoacetamide (IAA) in the dark for 45 min at room temperature. Proteins were then digested overnight at 37°C using sequencing-grade trypsin (1:50 enzyme-to-protein ratio). The digestion was quenched with formic acid, and peptides were desalted using C18 solid-phase extraction. The resulting peptides were analyzed by liquid chromatography-tandem mass spectrometry (LC-MS/MS) on a high-resolution mass spectrometer. Raw data were processed using Scaffold viewer, and protein identification was performed using the Uniprot database.

### Metal determinations by flame atomic absorbance spectroscopy

Three independent biological replicates of control and myoblasts KD for the SWI/SNF subunits were cultured under proliferating conditions and supplemented with 100 µM CuSO_4_ and 50 µM ZnSO_4_ as described above. Cells were washed with ice-cold PBS three times, scraped, and transferred to a 1.5 ml microcentrifuge tube. Samples were sonicated using a Bioruptor at medium intensity for 5 min with 30 s on-off cycles and total protein was quantified using the Bradford method (36). Samples were acid digested in concentrated HNO_3_ for ultra-trace analysis and diluted in purified water with a resistivity of 18 MΩ. Reagents and glassware were of analytical grade and cleaned with 3% HCl for 24 h to prevent contamination.

Comparative analysis of metal concentrations was carried out by triplicate measurements of Cu or Zn in each sample, using a flame atomic absorbance spectrophotometer (AAS; Agilent 55 AA Atomic Absorption Spectrometer) with Cu or Zn hollow cathode lamps as radiation source. Cu and Zn standard solutions (1000 mg/L; Sigma-Aldrich) were diluted as necessary to obtain working standards and determine the limits of detection and dynamic range of the method. Cu and Zn content on each sample was normalized to the initial cell mass as previously described (30, 31, 57–61).

### Visualization of labile Cu and Zn in C2C12 myoblasts using fluorescent sensors

To visualize labile Cu-pools in proliferating myoblasts we used the membrane-permeable fluorescent dyes CS1 for Cu^+^ and CD649.2 to detect Cu^2+^ following standard procedures (33, 62–65). Control and the SWI/SNF KD myoblasts were seeded in CellView Plates and treated with or without 100 µM CuSO_4_ as described. Before Cu probe incubation, cells were washed with PBS containing GlutaMax and CaCl_2_, then incubated with the probes (5 µM) in the dark for 10 min. Imaging was done using a Leica SP8 confocal microscope and analyzed with the Leica Application Suite X. The CS1 probe visualized Cu^+^ under 543 nm excitation and 566 nm emission and the CD649.2 probe for Cu^2+^ was imaged at 633 nm excitation, and fluorescence emission was collected in the range of 650-750 nm (62–64).

To detect free Zn^2+^ in proliferating C2C12 myoblasts, the cells were loaded with 2 μM Fluo-Zin3 AM cell permeant dye (Invitrogen, cat. No: F24195) in the corresponding culture media for 40 min at 37°C, as previously described (61). Cells were then washed with fresh media and allowed to rest for another 20 min at 37°C. Culture media was removed, the cells were washed with fresh PBS and immediately visualized using a Leica SP8 microscope. Images were analyzed with the Leica Application Suite X. Fluo-Zin3, AM has an excitation and emission maximum of 494/516 nm.

### Statistical Analysis

All statistical analysis was performed in Kaleidagraph (Version 4.1) or Microsoft Excel. Statistical significance was determined using a Student’s T-test. Any experiments where P < 0.05 were considered statistically significant.

### Models and diagrams

All drawings and models representing biological processes were created with BioRender.com.

## DATA AVAILABILITY

Genomic data sets have been deposited in the Gene Expression Omnibus (GEO) accession no. GSE253379.

## RESULTS

### *Baf180* knockdown myoblasts are sensitive to metal stress, while Cu and Zn supplementation rescued the proliferation defect observed in *Baf250a* and *Brd9* deficient cells

We previously demonstrated that *Baf250a* KD significantly reduced myoblast proliferation, whereas *Brd9* KD caused a modest yet notable decrease (12). In contrast, *Baf180* KD had no observable effect on proliferation compared to control cells (12). However, RNA-seq analyses revealed substantial alterations in the expression of genes involved in homeostasis maintenance in *Baf180* KD cells (12). To further investigate the role of SWI/SNF subunits in cellular stress responses, we assessed the sensitivity of C2C12 myoblasts, stably transduced with shRNA targeting *Baf180, Baf250a,* and *Brd9*, to metal stress. Successful KD at the protein level was confirmed by western blot (**Fig. 1A-C**).

**Figure 1.**
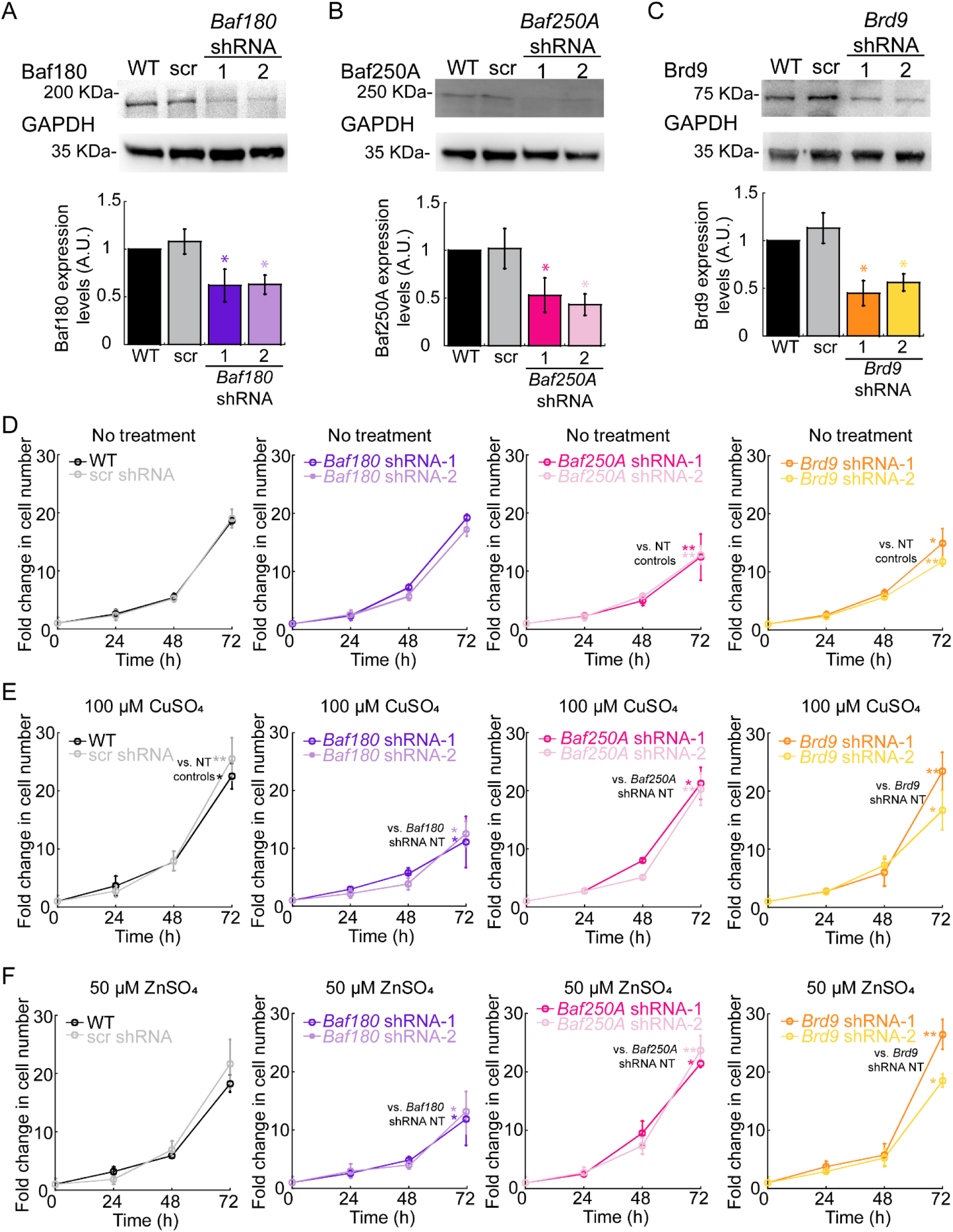
Expression and proliferation phenotype in response to metal supplementation of C2C12 myoblasts KD for *Baf180*, *Baf250a*, and *Brd9*. **(A-C)** Representative western blots (top) and quantification (bottom) of Baf180 **(A)**, Baf250a **(B)**, and Brd9 **(C)** protein levels in proliferating myoblasts, normalized to GAPDH as a loading control. Data represents the mean ± SE of three independent biological replicates. *P < 0.05 compared to scr control. **(D-F)** Proliferation analysis of WT, scr, and KD myoblasts over 72 h using cell counting assays for cells cultured under basal conditions **(D)** or supplemented with 100 µM CuSO_4_ **(E)** or 50 µM ZnSO_4_ **(F)**. *Baf180* KD myoblasts exhibits no significant difference under basal conditions, non-treated (NT), but this cell line sensitive to both metals. *Baf250* KD and Brd9 KD show reduced proliferation, which is rescued by metal supplementation. Data represent the mean ± SE of three independent experiments. *P < 0.05; **P < 0.01 compared to the same strain cultured in the absence of metals (NT).

To examine the impact of metal exposure on proliferative capacity, we cultured control and KD cell lines with varying concentrations of CuSO_4_ and ZnSO_4_ (**Fig. 1D,E; Supp.** Fig. 1-4). Cell counting and Pax7 immunochemical staining, a marker of myoblast proliferation, showed that wild type (WT) and scrambled control (scr) myoblasts tolerated CuSO_4_ concentrations up to 100 µM, exhibiting enhanced proliferation, as previously reported (**Fig. 1E, Supp.** Fig. 1,2; (30)). However, *Baf180* KD myoblasts displayed a reduction in cell numbers at 50 µM CuSO_4_ and higher (**Fig. 1E, Supp. Fig. 1,2**). Interestingly, supplementation with 50-100 µM CuSO_4_ rescued the proliferation defects observed in *Baf250a* and *Brd9* KD myoblasts, whereas concentrations above 200 µM were lethal to all tested cell lines (**Fig. 1E, Supp. Fig. 1,2**). Similarly, immunohistochemistry and cell counting assays in ZnSO_4_-treated cells showed that WT and scr myoblasts maintained normal proliferation at 50 µM ZnSO_4_, while *Baf180* KD myoblasts exhibited reduced proliferation at this concentration (**Fig. 1F, Supp. Fig. 3,4**). Treatment with 50 µM ZnSO_4_ restored the proliferation defect of *Baf250a* and *Brd9* KD myoblasts to WT-like levels (**Fig. 1F, Supp. Fig. 3,4**). These findings suggested that *Baf180* KD myoblasts exhibit heightened sensitivity to metal stress, and CuSO_4_ and ZnSO_4_ supplementation rescues the proliferation defects in *Baf250a* and *Brd9* KD myoblasts. Considering this information, we selected concentrations of 100 µM CuSO_4_ and 50 µM ZnSO_4_ for further experiments, as these conditions elicited changes in proliferation without killing a substantial number of cells.

### Disruption of metal homeostasis in myoblasts with *Baf250a* and *Brd9* KD contributes to proliferation defects

We have demonstrated that proliferating and differentiating myoblasts undergo a dynamic flux of transition metals and associated metalloproteins, essential for their proper subcellular distribution and utilization depending on the cellular stage. During myogenesis, Cu levels increase in the nucleus and cytosol (30, 31, 33), while differentiation induces Zn efflux, followed by gradual Zn recovery as myotubes mature (61). Given the well-established detrimental effects of metal imbalance on cellular function (66–69), we investigated how SWI/SNF subunit KD affects metal homeostasis in proliferating myoblasts by quantifying Cu and Zn levels using atomic absorption spectroscopy (AAS). Our findings revealed that *Baf250a* KD myoblasts exhibit a significant reduction in both Cu and Zn compared to WT and scr controls (**Fig. 2A,B**). Similarly, *Brd9* KD myoblasts show a pronounced Zn deficiency, whereas *Baf180* KD myoblasts do not display significant alterations in total metal content (**Fig. 2A,B**). However, CuSO_4_ and ZnSO_4_ supplementation results in a substantial accumulation of Cu in *Baf180* KD myoblasts (**Fig. 2C,D**), aligning with their observed sensitivity to metal stress. This suggests a model in which the impaired expression of genes essential for metal homeostasis, as revealed by our RNA-seq analyses (12), disrupts cellular metal regulation and likely contributes to the proliferation defects observed in Baf180 KD myoblasts upon metal supplementation.

**Figure 2.**
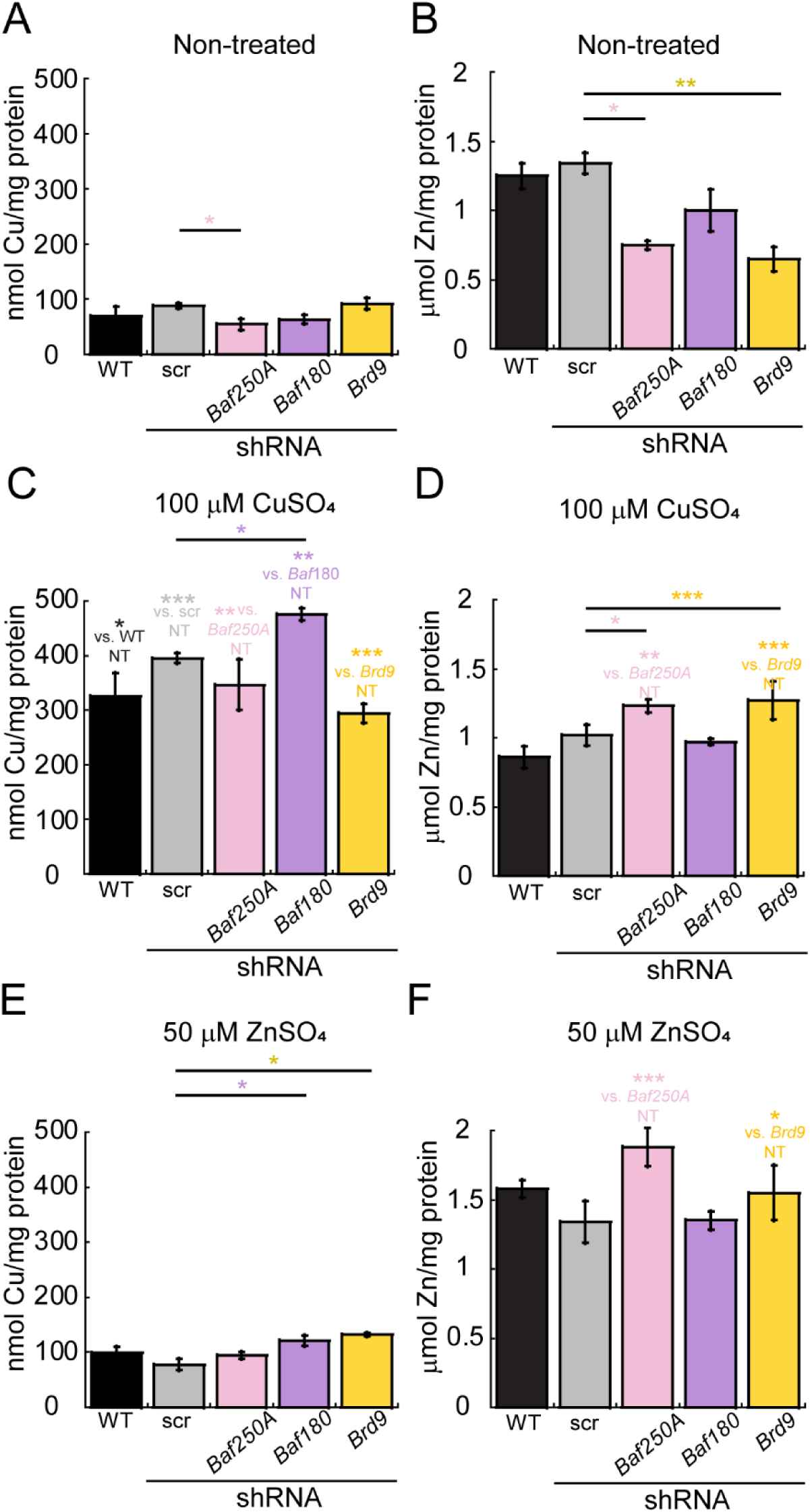
Knockdown of SWI/SNF subunits differentially disrupts Cu and Zn homeostasis in C2C12 myoblasts. Flame atomic absorption spectroscopy (AAS) analysis quantifying total intracellular Cu and Zn levels in C2C12 myoblasts under different conditions. **(A)** Quantification of total Cu and **(B)** total Zn for proliferating myoblasts cultured in basal medium without supplementation. **(C)** Quantification of total Cu and **(D)** total Zn for proliferating myoblasts cultured in the presence of 100 μM CuSO_4_. **(E)** Quantification of total Cu and **(F)** total Zn for proliferating myoblasts cultured with 50 μM ZnSO_4_. *Baf180* KD myoblasts exhibit significantly higher intracellular Cu accumulation upon CuSO_4_ supplementation, suggesting impaired Cu homeostasis. *Baf250A* KD have lower Cu and Zn content while *Brd9* KD myoblasts showed reduced levels of Zn. These defects are restored upon metal supplementation. Data represents the mean ± SE of three independent experiments. *P < 0.05; **P < 0.01; ***P < 0.001 compared to scr control.

Further AAS analysis revealed that Cu and Zn supplementation restores Cu levels in *Baf250a* KD myoblasts to those of control cells, while Zn deficiency in both *Baf250a* and *Brd9* KD myoblasts is corrected upon CuSO_4_ or ZnSO_4_ treatment (**Fig. 2E,F**). Interestingly, despite the significant Cu accumulation in *Baf180* KD cells upon supplementation, this excess Cu does not appear to be readily bioavailable for cellular processes, given that the rates of cell proliferation are reduced compared to no-treated cells.

To explore this possibility, we performed live-cell imaging using metal-specific fluorescent probes (**Fig. 3**; CS1 for Cu^+^, CD649.2 for Cu^2+^, and FluoZin3 for labile Zn^2+^). Confocal microscopy revealed that monovalent Cu was predominantly localized to subcellular compartments in non-treated control and *Brd9* KD cells (**Fig. 3A,B,E**), whereas *Baf180* (**Fig. 3C**) and *Baf250a* KD cells (**Fig. 3D**) displayed minimal labile Cu⁺ or Cu²⁺ signals. Notably, Cu supplementation rescued the labile Cu⁺ defect in *Baf250a* but not *Baf180* KD myoblasts (**Fig. 3C,D**), supporting our hypothesis that Cu in *Baf180* KD cells is trapped in a sequestered form rather than being bio available for cellular use. Zn supplementation elicited a similar response across all KD lines, further reinforcing the idea that myoblasts not only rely on these metals for proliferation but also utilize them to restore metal homeostasis.

**Figure 3.**
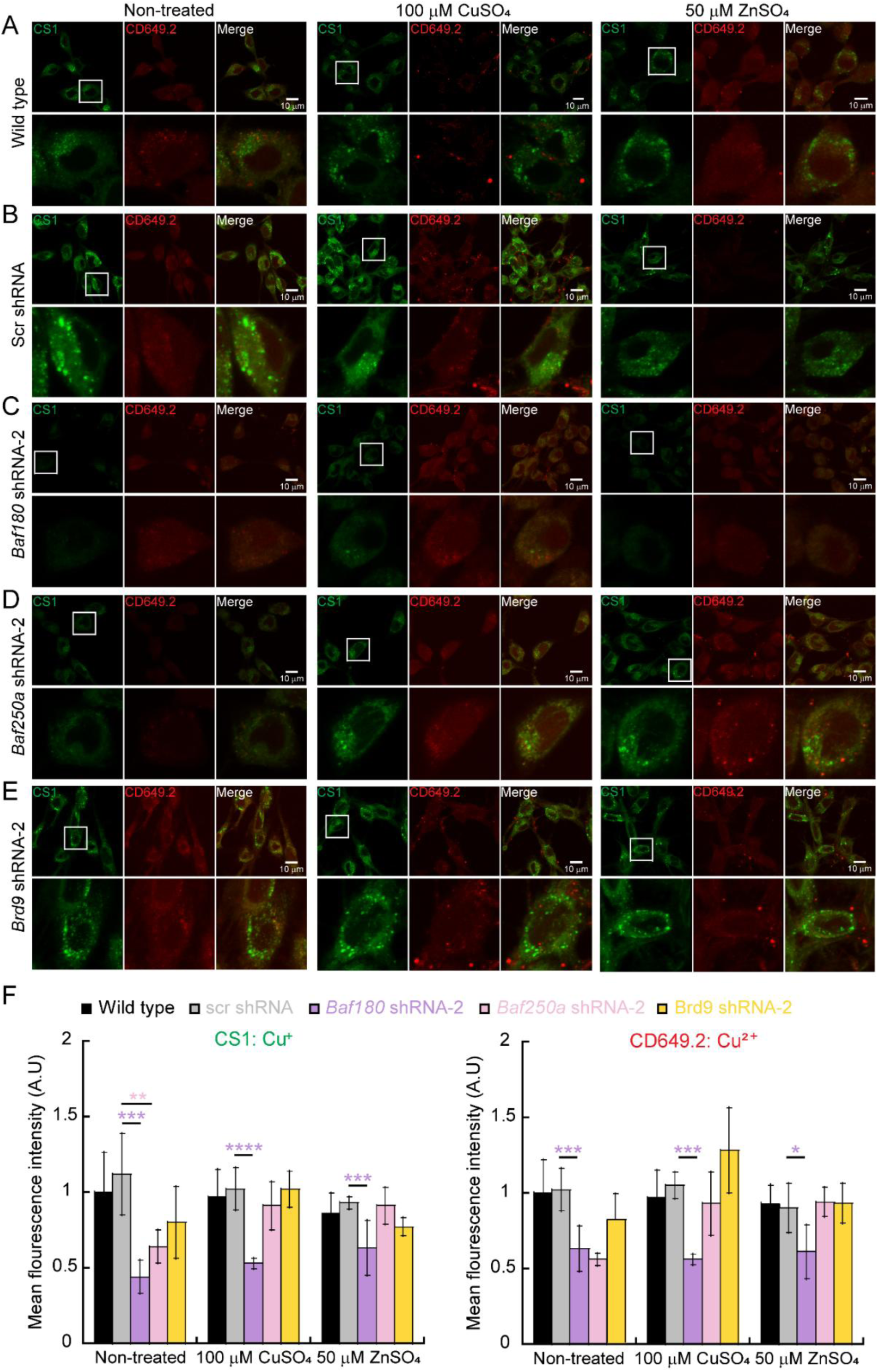
Distribution of labile Cu in control and SWI/SNF KD proliferating C2C12 myoblasts. Confocal microscopy live-cell analysis of labile Cu levels in wild-type **(A)**, Scr control **(B)**, and KD myoblasts for *Baf180* **(C)**, *Baf250a* **(D)**, and *Brd9* **(E)** under cultured for 48h in basal untreated media (NT) or supplemented with 100 μM CuSO_4_ or 50 μM ZnSO_4._ Labile Cu⁺ (green) was detected using CS1, while Cu²⁺ (red) was visualized using CD649.2. **F.** Quantification of the fluorescence of live-cell imaging for Cu^+^ with the CS1 probe (left panel) and Cu^2+^ with the CD649.2 probe (right panel) from proliferating C2C12 myoblasts. N = 3, *P < 0.05; ** P < 0.01, ***P < 0.001; ****P < 0.0001. *Baf180* KD myoblasts exhibited an decreased pools of labile Cu^+^ under all conditions tested, suggesting impaired copper mobilization. Scale bar: 10 μm.

Labile Zn imaging revealed cytosolic vesicular pools in proliferating control cells, as previously reported (**Supp.** Fig. 5 (61)), whereas all KD cell lines exhibited reduced labile Zn levels under basal conditions. An apparent reduction in labile Zn levels was detected in control cells, while Zn supplementation restored Zn pools in *Baf250a* and *Brd9* KD myoblasts but not in *Baf180* KD cells. The differential response of *Baf180* KD myoblasts suggests an intrinsic deficiency in metal regulatory mechanisms that prevents proper redistribution of both Cu and Zn. These findings highlight a critical role for SWI/SNF chromatin remodeling subunits in regulating metal homeostasis in myoblasts. Specifically, *Baf250a* and *Brd9* KD cells exhibit deficiencies in Cu and Zn that can be rescued by metal supplementation, whereas *Baf180* KD myoblasts accumulate excess Cu upon supplementation but fail to incorporate it into functional labile pools. This suggests that Baf180 is essential for maintaining the balance between metal uptake, utilization, and sequestration. The observed defects in proliferation and metal homeostasis in SWI/SNF KD cells suggest a broader role for chromatin remodeling complexes in coordinating metal-dependent cellular processes.

### Colocalization between BAF180 and MTF1 is higher than other SWI/SNF subunits upon metal treatment

So far, our data suggests a model where Baf180 is dispensable for myoblast proliferation and differentiation but may play a role in cellular stress responses, particularly metal homeostasis. These observations include our published RNA-seq analyses, which revealed downregulation of metal-regulating genes in *Baf180* KD myoblasts (12), and may partially explain the increased sensitivity to Cu and Zn supplementation-induced stress (**Fig. 1, Supp. Fig. 1-4**). Given the role of Mtf1 as a master regulator of metal homeostasis, we hypothesized a potential connection between Baf180 and Mtf1 in regulating cellular metal responses in proliferating C2C12 myoblasts. Thus, to investigate the potential interaction between the different SWI/SNF subunits and Mtf1, we assessed protein distribution and colocalization in WT myoblasts under untreated and metal-treated conditions. Confocal microscopy confirmed that Baf180 exhibited significantly higher colocalization with Mtf1 (**Fig. 4A,B,F**) than with Baf250a or Brd9 (**Fig. 4A; Supp. Fig. 6A,6B,7A,7B**) in control myoblasts. Further analysis of SWI/SNF subunit expression and colocalization for untreated, Cu-, or Zn-supplemented conditions revealed that Baf180 levels were significantly reduced in *Baf180* KD cells (**Fig. 4C**), while *Baf250a* (**Fig. 4D**) and *Brd9* KD (**Fig. 4E**) cells showed increased expression of Baf180 as nuclear speckles. Quantitative analyses showed a significant reduction in Baf180/Mtf1 colocalization in *Baf180* KD cells for all treatments (**Fig. 4F**), which can be explained by the reduced Baf180 expression in these cells (**Fig. 1; Supp. Fig. 8**). Despite the altered expression of Baf180 in KD cells, the expression of Mtf1 remained relatively unchanged. Notably, *Baf250a* and *Brd9* KD cells displayed enhanced Baf180/Mtf1 colocalization (**Fig. 4F**), possibly due to these nuclear speckles, the function of which remains unclear and warrants further investigation. Overall, quantification of the colocalization of Baf180 with Mtf1 in all cell lines revealed that this PBAF subunit exhibited the highest levels of colocalization among the tested SWI/SNF subunits (**Fig. 4F**). Colocalization levels remained relatively unchanged across untreated and metal-treated cells, indicating that while Baf180 has a strong interaction with Mtf1, this interaction does not appear to be influenced by metal supplementation. The overall higher colocalization of Baf180 with Mtf1 compared to other SWI/SNF subunits further supports the idea that Baf180 may play a unique role in metal homeostasis.

**Figure 4.**
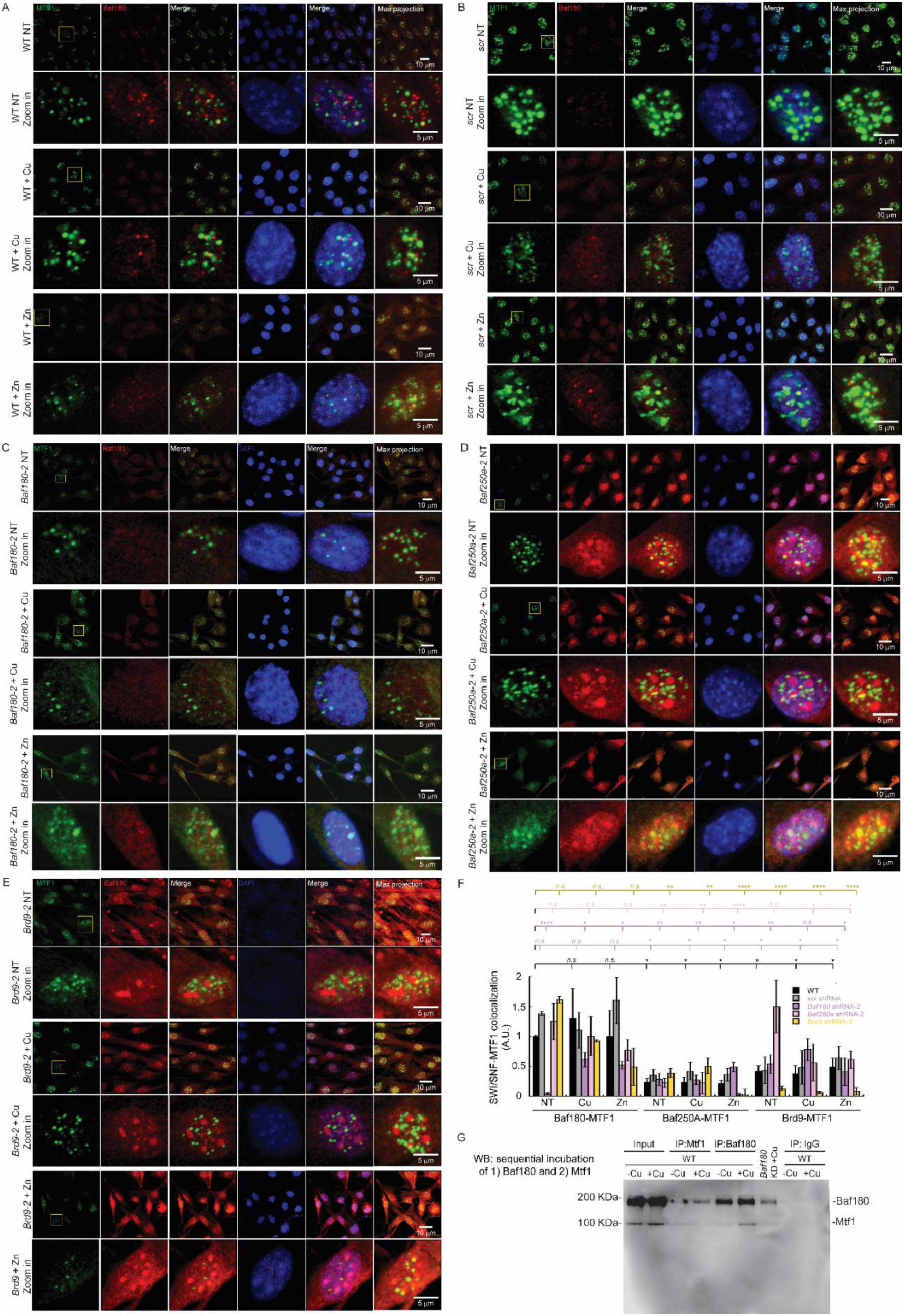
Baf180 and Mtf1 colocalize and interact in C2C12 control and SWI/SNF KD myoblasts upon metal treatment. Representative confocal microscopy images showing the expression and nuclear localization of Baf180 (red) and Mtf1 (green) in wild type **(A)**, scr **(B)**, *Baf180* KD **(C)**, *Baf250a* KD **(D)** and *Brd9* KD **(E)** C2C12 myoblasts. Nuclei are counterstained with DAPI (blue). **(F)** Quantification of Mtf1 colocalization with Baf180 in control and the different SWI/SNF KD myoblasts from panel A-E and from supplemental figures 6 and 7. Colocalization is disrupted in *Baf180* KD cells, while metal treatment modulates Mtf1-Baf180 interactions. Scale bars: 10 μm (panoramic views), 5 μm (zoomed-in images). Mean ± SE of three independent biological replicates. *P < 0.05, **P < 0.01, ****P < 0.0001. * Indicates significance relative to untreated WT cells. **(G)** Representative immunoprecipitation (IP) of Mtf1 or Baf180 in proliferating wild type myoblasts (N=3). Input, a Baf180 IP obtained from *Baf180* KD cells and an IgG IP were included as controls. Mtf1 exhibits significantly higher colocalization with Baf180 compared to Baf250a or Brd9 in WT proliferating myoblasts. To show both proteins, the western blot was developed using both the rabbit anti-Baf180 and the mouse anti-Mtf1 specific antibodies, incubated sequentially in the same membrane, followed by sequential incubation of species-specific secondary antibodies.

To confirm the direct interaction between Baf180 and Mtf1, we performed reciprocal immunoprecipitation (IP) assays using wild-type myoblasts cultured in the presence or absence of Cu. Mtf1 or Baf180 were immunoprecipitated with specific antibodies, followed by western blot analysis (Fig. 4G). Sequential probing for Baf180 and Mtf1 confirmed their interaction. As controls, we included Baf180 KD cells treated with Cu, immunoprecipitated using an anti-Baf180 antibody, and IgG as a negative control, ensuring the specificity of the interaction.

We also conducted unbiased IP experiments coupled to mass spectrometry of myoblasts supplemented with or without Cu (**Supp. Table 2**). This analysis identified Baf180, Baf250a, and Brg1 as interacting proteins, supporting our confocal microscopy findings and reinforcing the idea that MTF1 functions with these chromatin-remodeling complexes. This unbiased proteomic approach revealed a broader network of potential interactors, highlighting numerous candidates for further analysis. The identification of these factors suggests that MTF1 may play a key role in coordinating transcriptional responses to metal availability through interactions with chromatin regulators, warranting deeper investigation into the functional consequences of these associations.

### The interaction between Baf180 and Baf250a with Mtf1 is stronger than with Brd9

RNA-seq analyses indicated that Baf250a and Brd9 are not key regulators of cellular homeostasis. However, we compared their interactions with Mtf1 to that of Baf180.

Immunostaining and colocalization analyses showed minimal interaction between Baf250a and Mtf1 across all conditions (**Supp. Fig. 6**). Baf250a KD efficiency was confirmed by fluorescence analyses (**Supp. Fig. 8**), with Mtf1 expression remaining unchanged, suggesting no significant impact of Baf250a KD or metal treatments on its levels. Nonetheless, some colocalization was observed between Baf250a and Mtf1, corroborated by unbiased Mtf1 IP/MS (**Supp. Table 2**).

Brd9-Mtf1 interactions were also assessed via confocal microscopy (**Supp. Fig. 7**). Brd9 expression was unchanged in WT and Scr cells, with efficient KD confirmed by western blot (**Supp. Fig. 8**). Interestingly, *Baf180* KD cells treated with Cu exhibited increased Brd9 levels, suggesting a potential regulatory response to metal exposure. However, the increased Brd9/Mtf1 colocalization observed in *Baf250a* KD cells was likely due to high background interference, which consistently appeared in this cell line (**Supp. Fig. 7**). Together, these findings highlight a unique role for Baf180 in metal homeostasis, distinct from other SWI/SNF subunits. Confocal and IP experiments provided evidence for a direct interaction between Baf180 and Mtf1, reinforcing the hypothesis that Baf180 may facilitate metal homeostasis through Mtf1. While metal treatments did not significantly alter colocalization patterns, potential changes in Baf180/Mtf1 interaction under Zn treatment require further investigation. Overall, these results support the notion that Baf180, contributes to cellular metal regulation. Future studies should explore the functional consequences of this interaction, particularly in response to varying metal stress conditions, to fully elucidate the mechanisms by which Baf180 influences metal homeostasis in myoblasts.

### Transcriptional changes induced by metal supplementation in proliferating SWI/SNF-KD myoblasts

To understand the global impact of metal exposure on gene expression, we performed RNA-seq analysis on proliferating Scr control and *Baf180, Baf250a*, and *Brd9* KD C2C12 myoblasts treated with 100 µM CuSO_4_ or 50 µM ZnSO_4_. Principal component analysis (PCA) revealed distinct transcriptional signatures for each condition, with minimal variation between treated and untreated groups within the same KD, suggesting a stable global transcriptomic response to Cu and Zn exposure (**Supp. fig. 9**). Significant changes in gene expression were detected in non-treated KD cell lines, as previously reported (**Supp. Table 3,** (12)). Notably, *Baf250a* KD induced the most differentially expressed genes (DEGs) at 9362, followed by *Baf180* KD with 5968 DEGs (2967 upregulated and 3001 downregulated), while *Brd9* KD exhibited the fewest at 5345.

Analysis of metal-induced DEGs (**Supp. Tables 4,5**) revealed *Baf180* KD had the most changes (2818 with Cu, 461 with Zn), while *Baf250a* KD had 45 Cu- and 403 Zn-responsive genes. *Brd9* KD and Scr controls had minimal changes with Cu (20 DEGs), but Zn altered 1979 genes. Functional enrichment showed Cu in Scr cells upregulated oxidative stress genes (*Gsta4, Nqo1*) and repressed adhesion-related *Islr*. Zn treatment induced *Mt1* and *Slc30a1*, and repressed *Snora73b*. Cu also increased *Mmp12* (ECM remodeling) and repressed *Islr*, while Zn enhanced mitochondrial gene *mt-Nd5* and repressed *Scarna2*. GO terms for Cu-treated Scr cells highlighted oxidative stress and ECM remodeling; Zn treatment upregulated DNA/protein metabolism, cell cycle, and chromatin organization, and downregulated RNA processing and mitochondrial organization (**Supp. Fig. 10A,B**).

In *Baf180* KD cells (**Supp. Tables 4,5**), *Ralb* was most downregulated, while *Rassf2* and *Gas6* were upregulated. Cu treatment repressed *Synpo2l* and *Postn* (muscle regeneration), and induced *Klf4* (satellite cell differentiation). Zn suppressed *Ccdc8* (cell cycle) and induced *Car8* (pH homeostasis) and *Colec12* (immune response). GO analysis showed Cu downregulated mRNA metabolism, cell cycle, and DNA repair pathways, and upregulated *RUNX1* transcription, mitochondrial organization, and apoptosis. Zn downregulated neuron and heart development and DNA metabolism genes, while upregulating histone/DNA methylation and translation (**Fig. 5A,B**).

**Figure 5.**
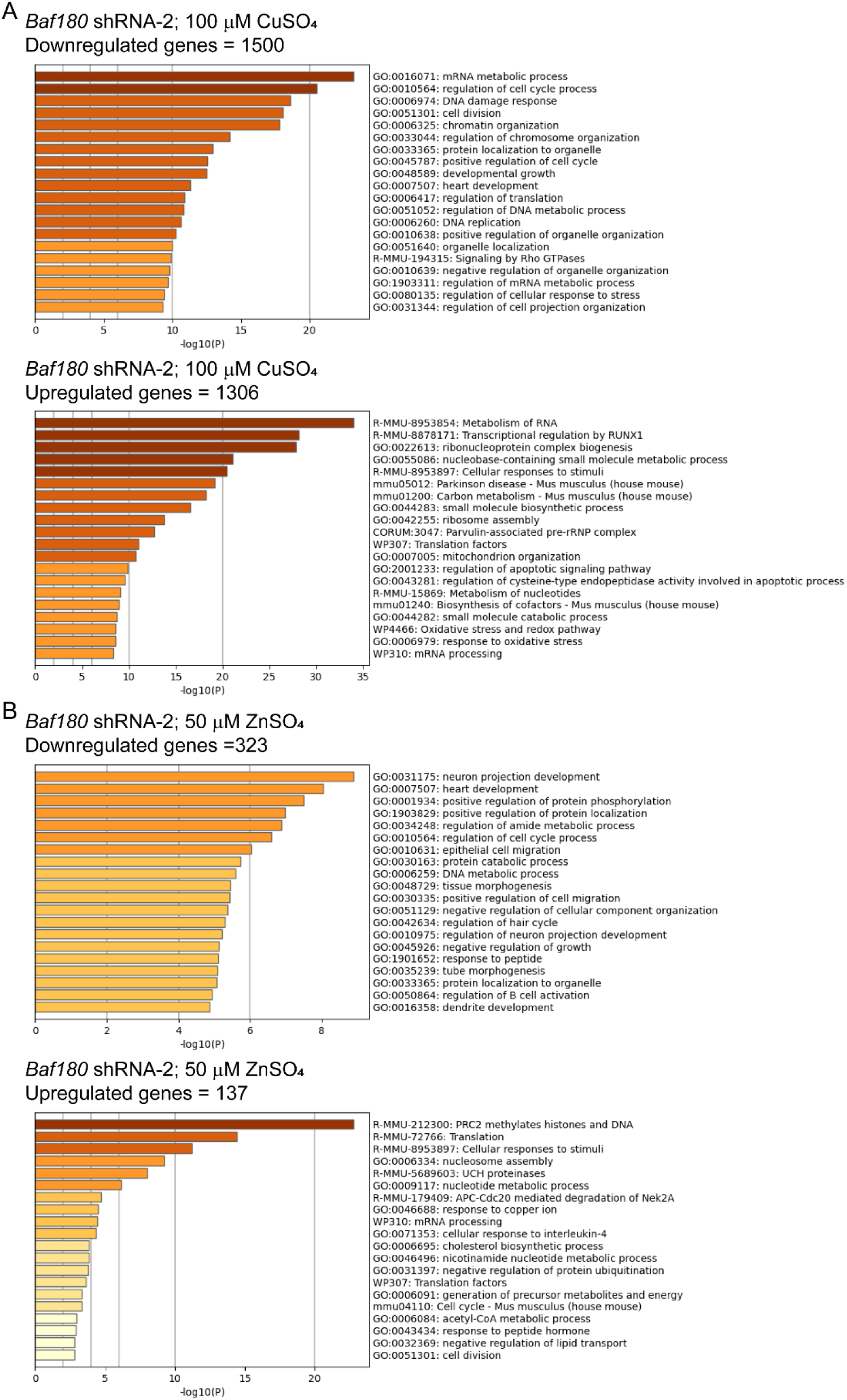
GO analysis of DEGs in *Baf180* KD myoblasts supplemented with metals. DEG identified from *Baf180* KD cells cultured in proliferation media supplemented with 100 μM CuSO_4_ **(A)** or 50 μM ZnSO_4_ (B) were compared to the same cell line cultured in the absence of metals (basal media). The patterns represent downregulation and upregulation of DEGs shown in **Supp. Table 4**.

In *Baf250a* KD cells, *Ubash3b* and *Cdk18* were downregulated, while *Mxra8* (myoblast fusion) and *Fgf10* (tissue repair) were upregulated. Cu caused few changes but repressed *Ethe1* (sulfide oxidation) and upregulated *Rab6b* (protein transport). Zn downregulated *Mfge8* (apoptotic clearance) and induced *Grem1* (BMP antagonist) and *Abca3* (xenobiotic resistance) (**Supp. Tables 4,5**). GO analysis showed Cu downregulated chemotaxis and response to stimuli, while upregulating oxidative stress and adhesion genes. Zn downregulated pluripotency, migration, and structure-related genes, and upregulated cell cycle, chromosome organization, and DNA replication genes. Stress response genes *Mt1* and *Slc30a1* were induced (**Supp.** Fig 11A,B).

*Brd9* KD cells had few DEGs with Cu. In untreated cells, *Selp* was most downregulated, while *Dkk3* (WNT regulator) and *Coro2a* (actin binding) were upregulated. Cu repressed *Ndrg4* and induced *Crlf1* (neuronal survival). Zn suppressed *Nrap* and *Synpo2l* (actin-binding) and induced *Ghr* (growth hormone receptor). Zn also induced *Mt1* and *Slc30a1*, indicating preserved stress responses. Several *Rny* noncoding RNAs involved in DNA replication were differentially regulated by Zn (**Supp. Table 4,5; Supp.** Fig 12A,B).

Common DEGs across conditions included *Gsta4* and *Nqo1* (oxidative stress); *Nqo1* was less induced in *Baf180* KD. Zn upregulated *Wnt4* in *Baf250a* and *Brd9* KDs, linked to myogenic proliferation. These findings suggest Cu and Zn influence genes tied to stress, proliferation, and homeostasis, highlighting their regulatory roles in *Baf250a* and *Brd9* KD myoblasts.

Notably, *Baf180* KD cells showed strong *Atp7a* downregulation after Cu treatment, suggesting a key regulatory effect on Cu export (**Supp. Table 5**). Zn also reduced *Atp7a* expression, though not significantly. In contrast, *Atp7a* levels in *Baf250a* and *Brd9* KD cells were unaffected by metal exposure. *Atp7b* expression was stable in *Baf250a* and *Baf180* KDs, but increased in *Brd9* KD with Zn. The Cu importer Ctr1 (*Slc31a1*) remained unchanged, indicating stable Cu uptake. These results suggest Cu homeostasis may be uniquely affected in *Baf180* KD myoblasts, with suppressed *Atp7a* indicating impaired Cu export and potential Cu toxicity.

### Mtf1 binding in proliferating myoblasts

Mtf1 is a transcription factor that activates gene expression in response to metal and oxidative stress (35, 70–75). It has been shown to bind myogenic genes in differentiating myoblasts, with Cu supplementation enhancing this binding and promoting myogenesis (31, 32). To examine the role of Mtf1 upon Cu supplementation, CUT&RUN was performed in proliferating WT C2C12 myoblasts treated with 100 μM CuSO_4_ for 48 h (**Supp. Table 6**). Mtf1 binding was measured using specific antibodies, with IgG as a control. As previously observed (31), Cu significantly increased Mtf1 binding at transcription start sites (TSS; **Fig. 6A,B**).

**Figure 6.**
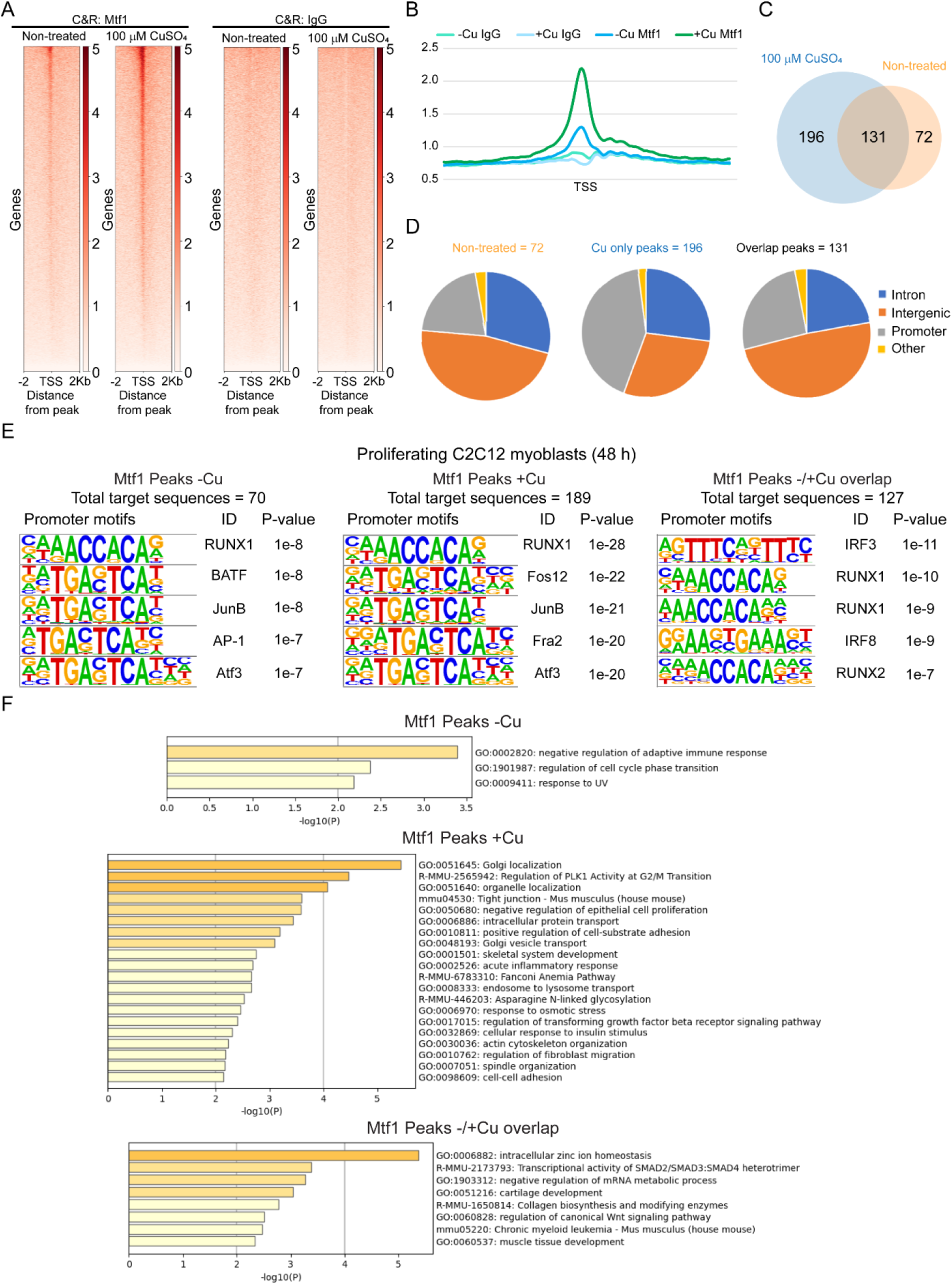
Cu supplementation enhances Mtf1 chromatin binding in proliferating myoblasts. CUT&RUN analysis illustrating Mtf1 binding dynamics in proliferating myoblasts under untreated conditions and following 100 μM CuSO_4_ supplementation for 48 h. **(A)** Combined CUT&RUN heatmaps from peak calling of three independent biological sequences obtained for each condition. Normal IgG used as control. **(B)** Aggregation plots of Mtf1 CUT&RUN data showing occupancy of Mtf1 and IgG over TSS’s in the presence or absence of Cu. **(C)** Overlap of CUT&RUN peaks of Mtf1 across the genome observed in proliferating cells in the presence or absence of Cu. See complete set of genes in **Supp. Table 5**. **(D)** Pie charts showing the binding site locations for Mtf1 in myoblasts cultured in the presence or absence of Cu and combined in both conditions. **(E)** Main changes of Mtf1 motif-binding dependent on Cu supplementation in proliferating C2C12 myoblasts myoblasts. Novel consensus DNA-binding motifs identified from peak calling for CUT&RUN within Mtf1 peaks in proliferating cells supplemented or not with Cu and in both conditions combined. The top five most significant motifs enriched, including the DNA logo, its corresponding transcription factor, and its P value are shown. **(F)** GO analysis of CUT&RUN data showing the main categories of genes that Mtf1 binds to in proliferating myoblasts in both the absence or presence of 100 μM CuSO_4_ and in both conditions combined. Cut-off was set at 2.0 of the -log(adjusted P value). Consensus DNA-binding motifs within Mtf1 peaks reveal distinct sequence preferences in Cu-treated versus untreated cells.

Genome-wide analysis identified 399 promoters bound by Mtf1: 196 in Cu-treated, 72 in untreated, and 131 in both conditions (**Fig. 6C; Supp. Table 6**). In untreated cells, Mtf1 binding was less enriched at promoters and more common in intergenic regions (**Fig. 6D**), suggesting Cu enhances promoter-specific binding and transcriptional activation.

Mtf1 target genes are involved in DNA repair, metabolism, and stress response (**Supp. Table 6**). Given our RNA-seq findings linking PBAF and Baf180 to DNA repair, a cooperative role with Mtf1 in genome maintenance is suggested. HOMER motif analysis identified Mtf1-associated transcription factor binding sites (**Fig. 6E**), including IRFs and Runx family motifs in both conditions. IRFs regulate immune responses (76), and *Runx1* supports myoblast proliferation (77). In Cu-treated cells, Mtf1 also bound *Fosl2*, *JunB*, and *ATF3* motifs— transcriptional activators including AP-1 factors that promote myogenesis via MyoD1 (78). These results support a role for Cu-activated Mtf1 in driving muscle proliferation through co-regulation with myogenic activators.

GO analysis confirmed Mtf1 binding to genes linked to homeostasis, stress response, and detoxification, consistent with its metal-regulatory function (**Fig. 6F; Supp. Table 6**). In untreated myoblasts, Mtf1 targeted genes involved in immune response, cell cycle, and UV stress response. Cu-specific targets included genes related to organelle/protein localization and skeletal muscle development. *PLK1* regulation, which is critical for mitosis and cell cycle progression (79), was among the top Cu-responsive categories, suggesting a role for Mtf1 in Cu-induced proliferation. Shared targets across conditions included genes involved in homeostasis, transcriptional regulation, collagen synthesis, *Wnt* signaling, and muscle development (**Fig. 6F**). Overall, CUT&RUN results demonstrate that Cu enhances Mtf1 binding at promoters, providing insight into its role in metal-induced cell growth and differentiation. Further studies are needed to explore how Mtf1 integrates Cu signals with transcriptional networks governing myoblast proliferation.

### Interaction of Mtf1 with gene promoters and its correlation to changes in gene expression driven by KD of SWI/SNF subunits

We integrated RNA-seq analyses of SWI/SNF subunit KDs with Mtf1 CUT&RUN data to examine potential links between Mtf1 and chromatin remodelers in metal homeostasis. Analyses included RNA-seq from all KD strains vs. Scr controls under various metal treatments, compared with Mtf1 CUT&RUN data from Cu-treated cells (**Supp. Table 7**). A focused comparison of Cu-treated cell line RNA-seq with Cu-stimulated Mtf1 CUT&RUN was used to evaluate specific correlations between Mtf1 and each chromatin remodeler (**Supp. Table 8, Supp. Fig. 13**). We examined genes that were repressed, activated, or unchanged by Cu in each KD line and were Mtf1-bound in proliferating Cu-treated myoblasts (**Supp. Tables 7,8**). Across all merged conditions, the overlap of RNA-seq and CUT&RUN data was insufficient for GO analysis.

Given the observed interaction between Mtf1 and Baf180, we expected greater Mtf1 binding correlation in *Baf180* KD cells. In this KD line, Cu-induced genes included transcription and signaling regulators (*Map3k3, Mafk, Jun, Ncln, Nelfcd*), protein synthesis and mitochondrial function genes (*Mrpl45, Rps5, Rpl37, Mrpl18, Rps19*), and metabolic genes (*Got1, Eno1, Pgd, Pkm, Gsto1*). Membrane dynamics genes (*Cdc37, Plec*) and stress response regulators (*Parp3, Dcaf15, Dusp6*) were also upregulated. Downregulated genes included the lipid metabolism gene *Abhd17a*. Repressed transcription and RNA processing genes included *Snd1*, while *Fiz1* and *Ankrd2*, involved in signaling and cytoskeletal function, also showed decreased expression. Cu-independent genes included those involved in metabolism (*Ndufv3, Gstm1, Acaa1a, Sdhb*), RNA regulation (*Ppp1r14b, Khsrp, Snhg1*), cell structure (*Podnl1, Fkbp10, Lmna, Vim, Vcp*), and signaling (*Ppib, Htra2, Stub1, Eif5a*).

In *Baf250a* KD cells, Cu-upregulated Mtf1 targets included *Pgd* and *Txnrd1*, linked to NADPH generation and redox balance, respectively. No Cu-repressed genes were identified, but *Gstm1* appeared Cu-independent, consistent with its detoxification role. In *Brd9* KD cells, Cu also induced *Pgd* and *Txnrd1*, and upregulated *Hmga2*, a chromatin-associated protein influencing proliferation and differentiation. No Cu-repressed genes were identified, but metal homeostasis genes *Mt1, Mt2*, and detoxification gene *Gstm1* were Cu-independent Mtf1 targets.

These findings reveal distinct Cu-responsive gene programs linked to Mtf1 activity and suggest cooperation with the SWI/SNF complex, particularly the PBAF subfamily. Genes varied in their responses—some activated, repressed, or unaffected—highlighting the complexity of Cu-regulated transcription and the need for further investigation into alternative regulatory pathways.

## DISCUSSION

Mammalian SWI/SNF complexes are ATP-dependent chromatin remodelers that reposition nucleosomes to regulate transcription (6, 7). They fall into three classes: BAF, PBAF, and ncBAF (11), each with unique subunits suggesting distinct roles. Our prior work identified BAF as a key regulator of myoblast proliferation via Pax7 expression (12), and essential for differentiation by activating myogenic genes (13). While ncBAF indirectly influenced myogenesis (12, 13), PBAF showed little effect on proliferation or differentiation, but RNA-seq from this study indicated a possible role in metal regulation (12).

### Role of SWI/SNF complexes in metal homeostasis and myoblast proliferation

We further examined SWI/SNF roles in metal regulation and homeostasis in proliferating myoblasts. Baf180 (PBAF complex) was essential for Cu and Zn homeostasis, as its KD impaired metal handling and inhibited myoblast proliferation. Unlike *Baf250a* and *Brd9* KDs, which showed no metal-stress proliferation defects, *Baf180* KD myoblasts were uniquely sensitive. These cells accumulated intracellular Cu but lacked labile Cu⁺, suggesting impaired Cu mobilization for cuproprotein function. RNA-seq revealed downregulation of *Atp7a*, a Cu exporter, in *Baf180* KD cells under Cu stress, which likely contributed to Cu accumulation in an unutilized, non-bioavailable state, resulting in cellular toxicity and impaired proliferation. *Mt1* and *Gsta4*, genes for metal sequestration and oxidative stress defense, were also not properly induced, highlighting a compromised stress response and contributing to proliferation failure by potentially making these cells more vulnerable to metal-induced toxicity. Sequence data suggested further impairment in DNA damage and metal stress responses in *Baf180*-deficient cells.

Previously, we showed that *Baf250a* and *Brd9* KDs impair proliferation (12), but Cu or Zn supplementation rescued this defect. Our current data show these subunits are not essential for metal response. Wnt4 emerged as a potential mediator, as its expression increased with Cu in both KD lines. *Wnt4*, key in myoblast proliferation and differentiation, enhances fast-type muscle development and upregulates *Pax7* and *MyoD1* expression in chick embryos, increasing muscle mass and fast MyHC expression (80). Wnt4 was absent in proliferating cells but induced during differentiation in C2C12 and satellite cells (81, 82), suggesting a differentiation-promoting role via β-catenin signaling (81). Interestingly, Pax7 expression was restored by Zn only in *Baf250a* KD, not *Brd9* KD cells, suggesting different mechanisms.

We also explored Cu and Zn in epigenetic regulation, as these metals can influence chromatin remodelers and transcription (27, 31). Zn is a cofactor in histone-modifying enzymes (83), and both metals regulate HDACs and DNMTs. Since Baf250a and Brd9 are part of SWI/SNF, their loss may alter chromatin accessibility. Zn may compensate by restoring chromatin dynamics. Cu and Zn also induced *Mt1* and *Gsta4*, possibly replacing stress genes lost in SWI/SNF KDs. As cofactors for SOD1/3, they support antioxidant defenses, improving cell growth. They also modulate cell cycle proteins, including p53 (67, 84–88), potentially rescuing proliferation. Further work is needed to detail how metals directly affect transcription in these KD cells.

### Interaction between Baf180 and Mtf1 in metal homeostasis

At the molecular level, Baf180 interacted with Mtf1, a transcription factor central to metal response. Immunofluorescence showed nuclear colocalization, and IP assays confirmed interaction. Mtf1 binding to chromatin increased with Cu, but in *Baf180* KD cells, its expression was downregulated under Cu stress. This suggests Baf180 may stabilize Mtf1 or aid its chromatin recruitment. CUT&RUN analysis found Mtf1 bound to *Runx* motifs—linked to myogenesis and stress adaptation—suggesting Baf180 may help Mtf1 regulate both metal response and muscle proliferation.

### The PBAF Complex in Cellular Stress and Homeostasis

Baf180 also supports stress adaptation beyond myoblasts. In hematopoietic stem cells, *Baf180* loss causes cell cycle arrest (23), and in *Caenorhabditis elegans*, it increases thermal stress sensitivity (89). In humans, Baf180 is frequently mutated in renal carcinoma (89), due in part to p53-mediated degradation (90), impairing stress responses and promoting tumor growth. Together with our findings, this supports Baf180’s broad role in environmental stress responses, especially metal homeostasis. Its loss disrupts Cu/Zn regulation and Mtf1 interaction, impairing proliferation. These results position PBAF as a key regulator of myogenesis and suggest targeting metal homeostasis may benefit muscle-related conditions which may result from dysregulation of these micronutrients.

## CONCLUSION

We propose that Baf180 interacts with Mtf1 to regulate metal-responsive gene expression, likely through chromatin remodeling by PBAF. This interaction is essential for Mtf1 stability and recruitment to chromatin under metal stress. CUT&RUN analysis further revealed that Mtf1 binding sites are enriched for Runx motifs, linking Baf180 to transcriptional programs governing both myogenesis and stress adaptation. These findings suggest that Baf180 plays a critical role in maintaining myoblast proliferation by coordinating metal homeostasis and transcriptional responses to environmental stressors. Our work also revealed that that Cu and Zn supplementation provides a compensatory mechanism that partially restores proliferation in *Baf250a* and *Brd9* KD myoblasts. This rescue effect likely involves a combination of chromatin remodeling, transcriptional activation, oxidative stress mitigation, and direct regulation of cell cycle factors. Further studies are needed to delineate the precise molecular pathways through which Cu and Zn exert their effects, which could provide valuable insights into metal ion homeostasis in muscle biology and disease contexts.

## ACKNOWLEGEMENTS

The authors are thankful to Dr. Rafael Pimenta for his technical assistance. This research was supported in part by the University of Pittsburgh Center for Research Computing, RRID:SCR_022735, through the resources provided. Specifically, this work used the HTC cluster, which is supported by NIH award number S10OD028483. This project used the University of Pittsburgh HSCRF Genomics Research Core, RRID: SCR_018301 NGS sequencing services, with special thanks to the Assistant Director, Will MacDonald. This work was supported by the National Institutes of Health [R35 GM133732 to S.J.H.]

## AUTHOR CONTRIBUTIONS

**Conceptualization:** Teresita Padilla-Benavides.

**Data curation:** Nick Carulli, Emma E. Johnston, David C. Klein, Odette Verdejo-Torres, Anand Parikh, Antonio Rivera, Michael Quinteros, Sarah J. Hainer

**Formal analysis:** Nick Carulli, Emma E. Johnston, David C. Klein, Odette Verdejo-Torres, Anand Parikh, Antonio Rivera, Michael Quinteros.

**Funding acquisition:** Christopher J. Chang, Sarah J. Hainer, Teresita Padilla-Benavides.

**Investigation:** Nick Carulli, Emma E. Johnston, David C. Klein, Odette Verdejo-Torres, Anand Parikh, Antonio Rivera, Michael Quinteros, Aidan T. Pezacki, Teresita Padilla-Benavides.

**Methodology:** David C. Klein, Michael Quinteros, Aidan T. Pezacki, Teresita Padilla-Benavides.

**Project administration:** Teresita Padilla-Benavides.

**Resources:** Christopher J. Chang, Sarah J. Hainer, Teresita Padilla-Benavides.

**Software:** Anand Parikh, David C. Klein, Sarah J. Hainer, Teresita Padilla-Benavides.

**Supervision:** Sarah J. Hainer, Teresita Padilla-Benavides.

**Validation:** Nick Carulli, Emma E. Johnston, Odette Verdejo-Torres, Antonio Rivera, Michael Quinteros.

**Visualization:** Nick Carulli, Emma E. Johnston, David C. Klein, Anand Parikh, Teresita Padilla-Benavides.

**Writing – original draft:** Nick Carulli, Emma E. Johnston, Teresita Padilla-Benavides.

**Writing – review & editing:** Nick Carulli, Emma E. Johnston, David C. Klein, Odette Verdejo-Torres, Anand Parikh, Antonio Rivera, Michael Quinteros, Aidan T. Pezacki, Christopher J. Chang, Sarah J. Hainer, Teresita Padilla-Benavides.

