## Supplementary material for "The PBAF chromatin remodeling complex contributes to metal homeostasis through Mtf1 regulation": Supp. figures and supp tables 1&4

[2] Department of Biological Sciences. University of Pittsburgh, Pittsburgh, PA. USA

[3] Department of Chemistry, University of California, Berkeley, CA 94720. USA

[4] Department of Chemistry, Princeton University, Princeton, NJ, USA. USA

[5] Department of Molecular and Cell Biology, University of California, Berkeley, CA 94720. USA

[6] Helen Wills Neuroscience Institute, University of California, Berkeley, CA 94720. USA

[#] Current affiliation: Tisch Multiple Sclerosis Research Center of New York, New York, NY, 10019, USA

[€] Current affiliation: Merck & Co., 770 Sumneytown Pike, West Point, PA 19486

[&] Current affiliation: Department of Molecular, Cell and Cancer Biology, University of Massachusetts Medical School, Worcester, MA, 01605, USA

[@] Current affiliation: Department of Molecular Biology and Genetics, The Johns Hopkins University School of Medicine, Baltimore, MD, 21205, USA

[$] These authors contributed equally to this project.

### SUPPLEMENTAL MATERIALS

### TABLE OF CONTENTS

**SUPPLEMENTAL REFERENCES**

Supplemental Figure 1. Cu supplementation modulates myoblast proliferation in SWI/SNF KD cells.

Supplemental Figure 2. Zn restores proliferation in *Baf250a* and *Brd9* KD myoblasts but inhibits *Baf180* KD Cells.

Supplemental Figure 3. $CuSO_4$ supplementation impairs *Baf180* KD myoblast proliferation but restores the proliferation defect in *Baf250A* and *Brd9* KD myoblasts.

Supplemental Figure 4. $ZnSO_4$ supplementation impairs *Baf180* KD myoblast proliferation but restores the proliferation defect of *Baf250A* and *Brd9* KD myoblasts.

Supplemental Figure 5. Distribution of labile Zn in control and SWI/SNF KD proliferating C2C12 myoblasts.

Supplemental Figure 6. Expression and nuclear localization of Baf250a in proliferating C2C12 myoblasts.

Supplemental Figure 7: Expression and nuclear localization of Brd9 in proliferating C2C12 myoblasts.

Supplemental Figure 8. Quantification of SWI/SNF subunit protein levels in proliferating C2C12 myoblasts.

Supplemental Figure 9. Principal component analysis reveals consistent global transcriptome profiles across conditions.

Supplemental Figure 10. GO analysis of DEGs in scr control myoblasts supplemented with metals.

Supplemental Figure 11. GO analysis of DEGs in Baf250a KD myoblasts supplemented with metals.

Supplemental Figure 12. GO analysis of DEGs in Brd9 KD myoblasts supplemented with metals.

Supplemental Figure 13. MTF1 chromatin binding correlates with DEGs in SWI/SNF KD myoblasts under Cu exposure.

**SUPPLEMENTAL TABLES**

Supplemental Table 1. Plasmids used in this study

Supplemental table 2. MTF1 IP-MS data.

Supplemental table 3. RNAseq. DEG SCR vs BAF subunits KD +/- metals.

Supplemental table 4. Summary of DEG genes from metal-treated KD and scr control cells.

Supplemental table 5. RNAseq. DEG within BAF subunits KD strains +/- metals.

Supplemental Table 6. CUT&RUN. Mtf1 Annotated peaks and motifs. All conditions.

Supplemental Table 7. CUT&RUN – RNAseq integration. Mtf1 supplemented with Cu integrated with RNAseq comparison beteem similar metal treatment of KD vs SCR analyses.

Supplemental Table 8. CUT&RUN – RNAseq integration. Mtf1 supplemented with Cu integrated with RNAseq comparison of Cu treatment within strains analyses.

#### SUPPLEMENTAL REFERENCES

### SUPPLEMENTAL FIGURES

#### SUPPLEMENTAL FIGURE 1

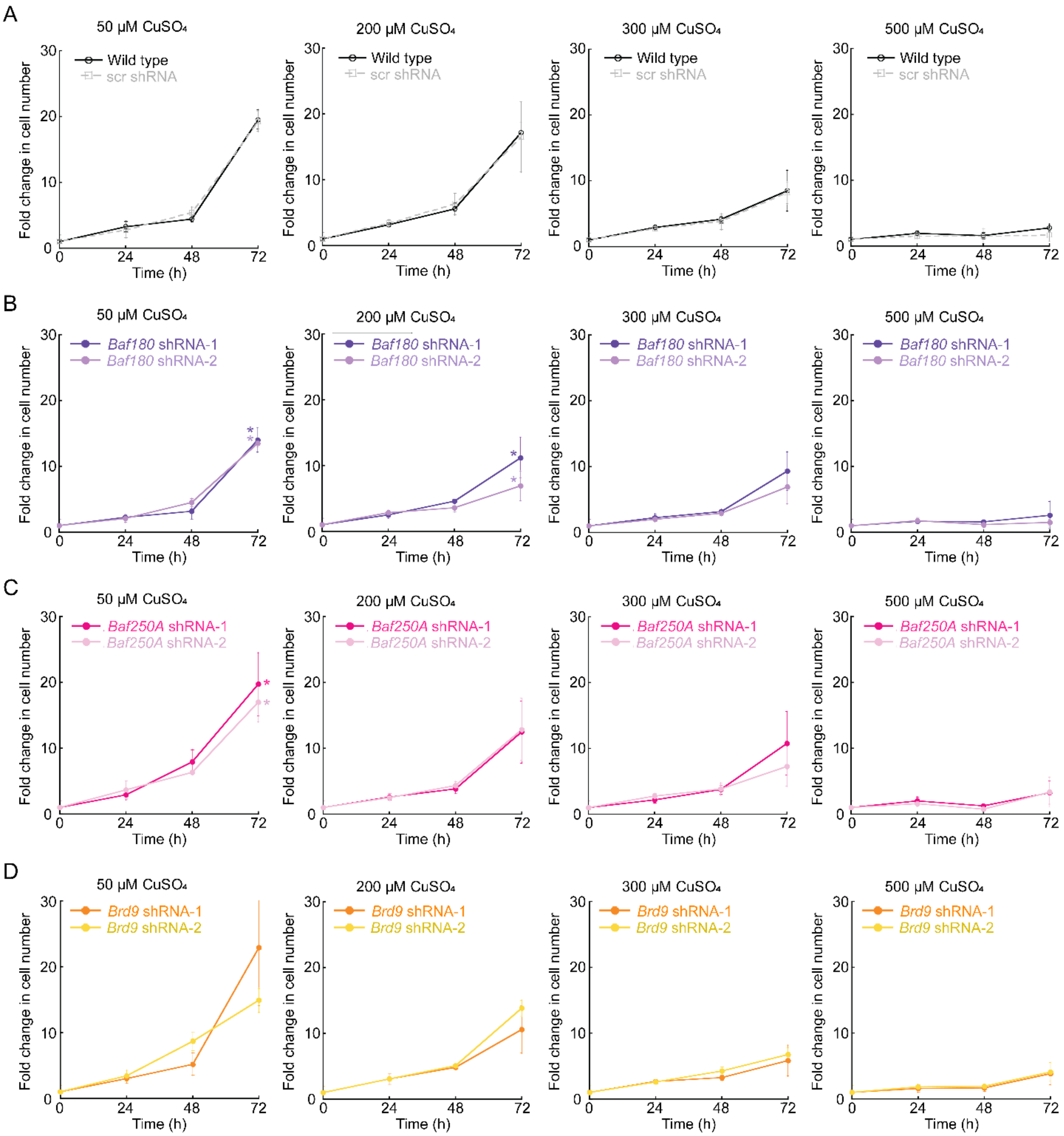


**Supplemental Figure 1. Cu supplementation modulates myoblast proliferation in SWI/SNF KD cells.** Cell counting assay of proliferating of C2C12 myoblasts treated with 50, 200, 300, and 500 µM $CuSO_4$ over 72 h. **(A)** Wild type and scrambled shRNA (scr) C2C12 myoblasts tolerate $CuSO_4$ concentrations up to 200 µM. **(B)** *Baf180* KD myoblasts exhibit reduced proliferation upon $CuSO_4$ treatment. **(C)** *Baf250a* KD myoblasts regain wild type-like proliferation with $CuSO_4$ supplementation. **(D)** *Brd9* KD myoblasts also recover wild type-like proliferation rates with $CuSO_4$ supplementation. See **Figure 1** for non-treated and 100 µM $CuSO_4$ concentrations, as these were the selected conditions used in this paper. The results indicate that Cu availability differentially affects myoblast proliferation, inhibiting *Baf180*-deficient cells while restoring proliferation in *Baf250a* and *Brd9* KD myoblasts. Data represents the mean ± SE of three independent experiments. *$P < 0.05$ compared to the same strain cultured in the absence of metals (NT).

**SUPPLEMENTAL FIGURE 2**

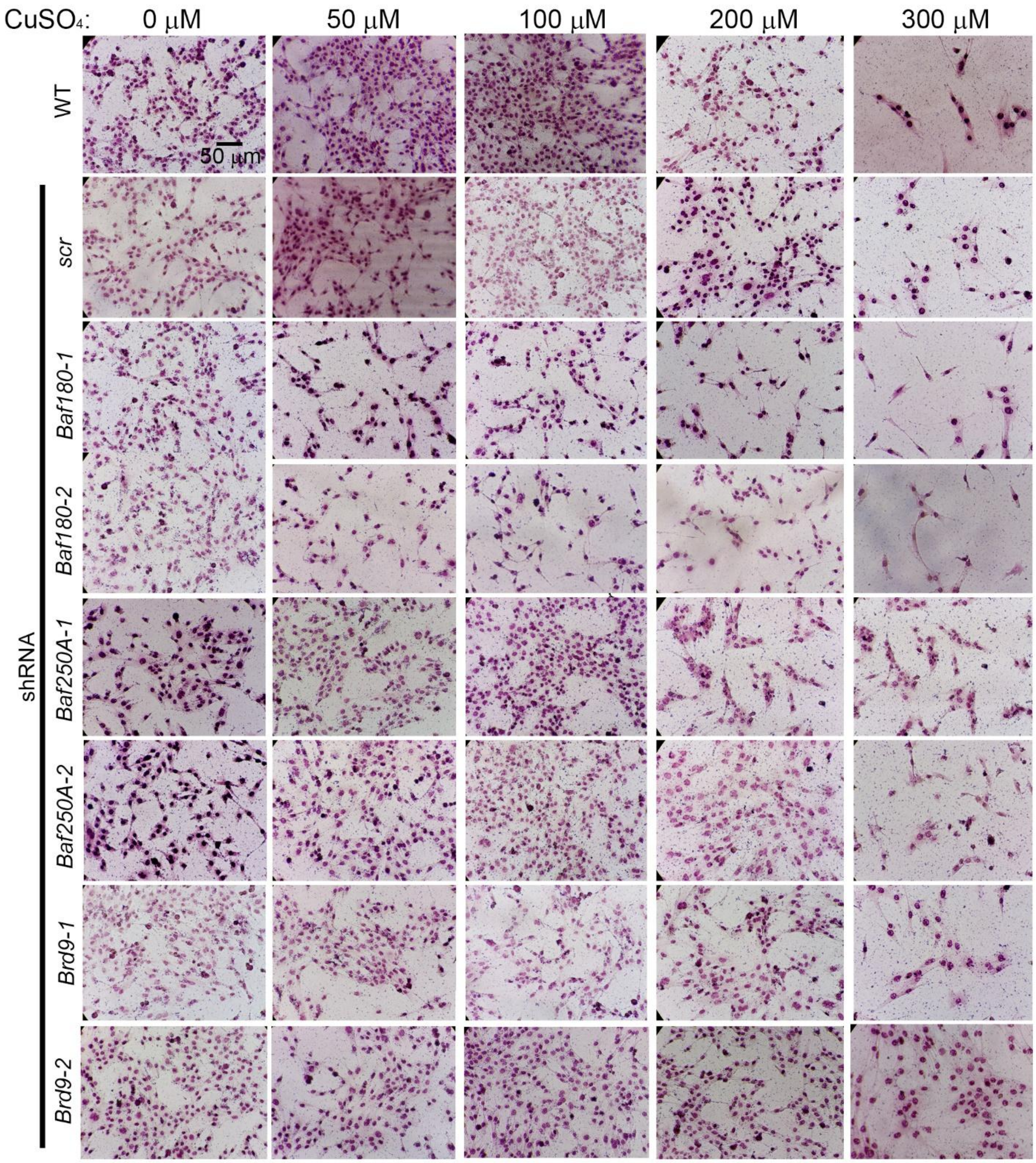


**Supplemental Figure 2. $CuSO_4$ supplementation impairs *Baf180* KD myoblast proliferation but restores the proliferation defect in *Baf250A* and *Brd9* KD myoblasts.** Representative light micrographs of proliferating WT, scr, *Baf180* KD*, Baf250A* KD*,* or *Brd9* KD C2C12 myoblasts supplemented with 0, 50, 100, 200, and 300 μM CuSO4 over 48 h and immunostained for Pax7.

**SUPPLEMENTAL FIGURE 3**

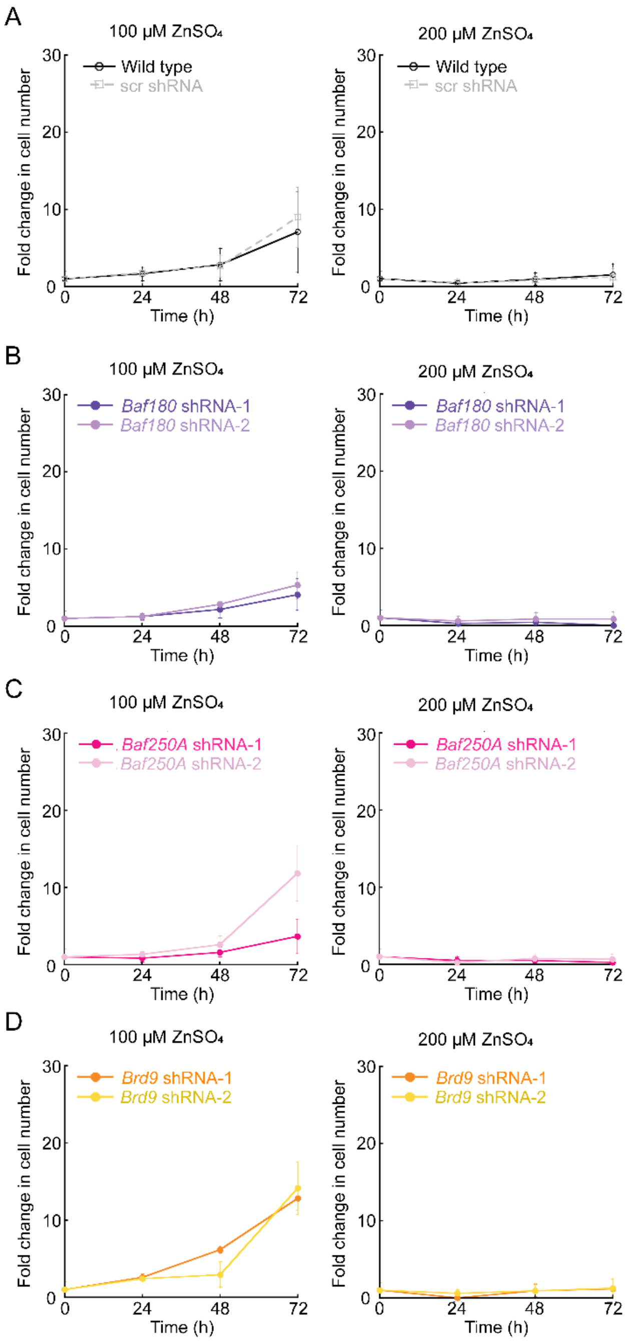


**Supplemental Figure 3. Zn restores proliferation in *Baf250a* and *Brd9* KD myoblasts but inhibits *Baf180* KD Cells.** Cell counting assay of proliferating of C2C12 myoblasts treated with 100 and 200 µM $ZnSO_4$ over 72 h. **(A)** Wild type and scrambled shRNA (scr) C2C12 myoblasts are sensitive to $ZnSO_4$ over 50 µM as previously shown (1). *Baf180* **(B)** and *Baf250a* **(C)** KD myoblasts exhibit reduced proliferation upon $ZnSO_4$ treatment. **(D)** *Brd9* KD myoblasts also recover wild type-like proliferation rates with $ZnSO_4$ supplementation. See **Figure 1** for non-treated and 50 µM $ZnSO_4$ concentrations, as these were the selected conditions used in this paper. These data suggest that Zn supplementation restores proliferation defects in *Baf250a*- and *Brd9*-deficient myoblasts at low concentrations and negatively impacts *Baf180* KD cells, highlighting distinct roles for SWI/SNF subunits in metal homeostasis and muscle cell growth. Data represents the mean ± SE of three independent experiments.

**SUPPLEMENTAL FIGURE 4**

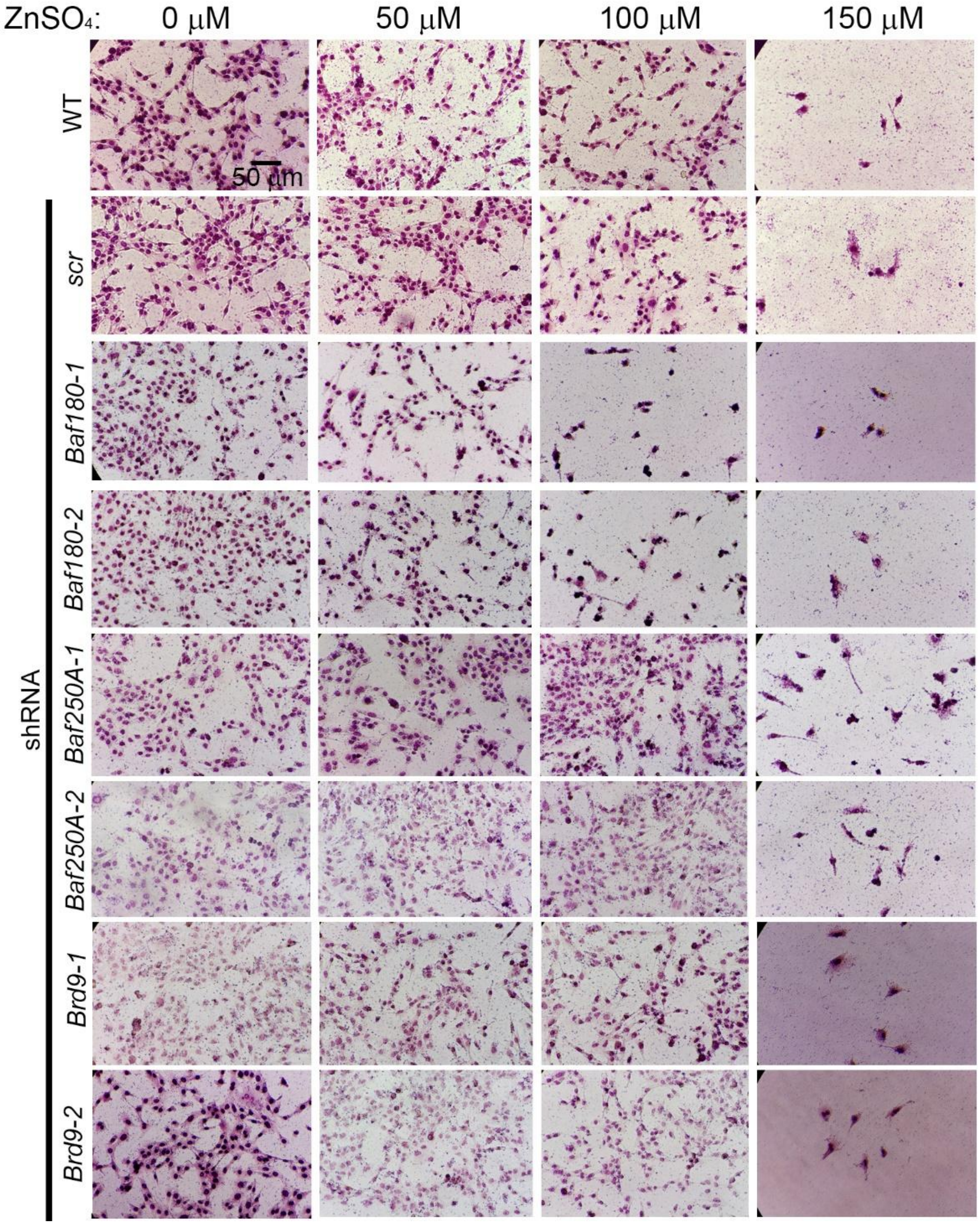


**Supplemental Figure 4. $ZnSO_4$ supplementation impairs *Baf180* KD myoblast proliferation but restores the proliferation defect of *Baf250A* and *Brd9* KD myoblasts.** Representative light micrographs of proliferating WT, scr, *Baf180* KD*, Baf250A* KD*,* or *Brd9* KD C2C12 myoblasts supplemented with 0, 50, 100, and 150 µM $ZnSO_4$ over 48 h and immunostained for Pax7.

#### SUPPLEMENTAL FIGURE 5

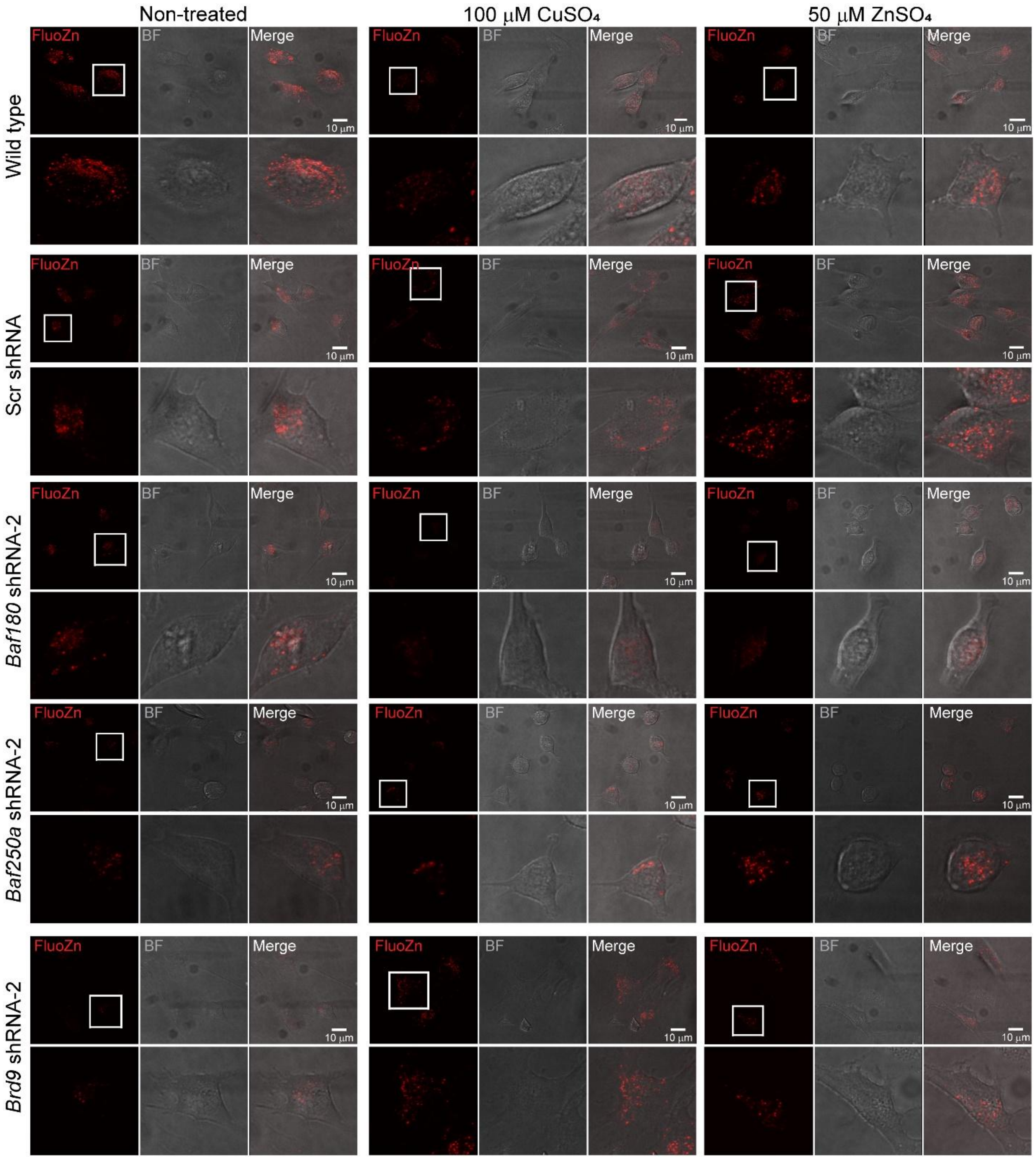


**Supplemental Figure 5. Distribution of labile Zn in control and SWI/SNF KD proliferating C2C12 myoblasts.** Confocal microscopy live-cell analysis of labile Zn in wild-type (WT), Scr control, and KD myoblasts for *Baf180*, *Baf250a*, and *Brd9* under cultured for 48 h in basal, untreated media (NT) or supplemented with 100 μM $CuSO_4$ or 50 μM $ZnSO_4$. Labile Zn (red) was detected using FluoZn3. Baf250a and Brd9 KD myoblasts exhibited a decreased signal of labile Zn when cells are cultured in basal media but recover upon not supplemented with metals. Scale bar: 10 μm.

#### SUPPLEMENTAL FIGURE 6

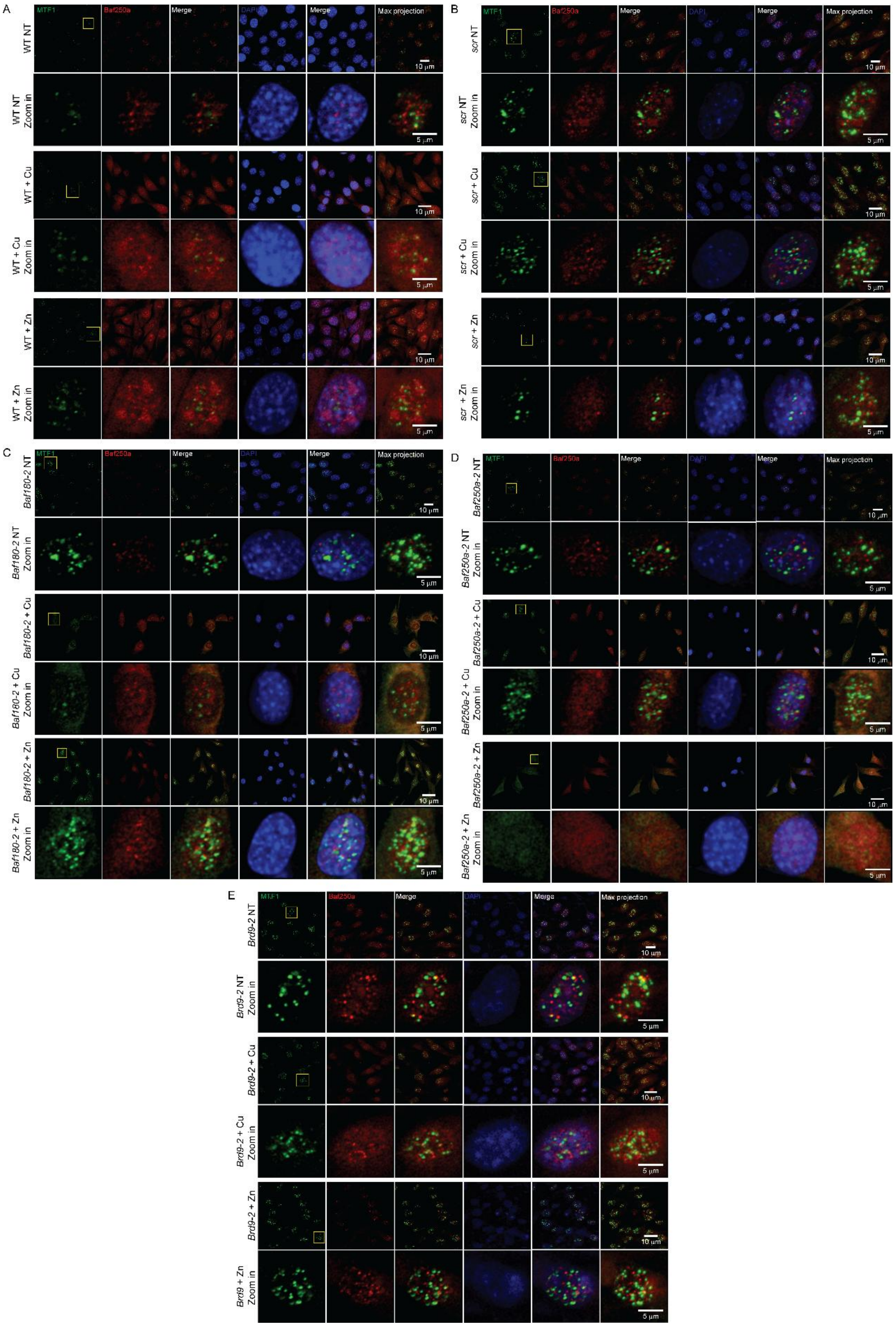


**Supplemental Figure 6. Expression and nuclear localization of Baf250a in proliferating C2C12 myoblasts.** Representative confocal microscopy images showing the expression and localization of Baf250a (red), Mtf1 (green), and DAPI (blue) in proliferating wild type (A), scr (B), *Baf180* KD (C), *Baf250a* KD (D) and *Brd9* KD (E) C2C12 myoblasts. Nuclei are counterstained with DAPI (blue). Non-treated myoblasts exhibit nuclear Baf250a localization, with limited colocalization observed between Baf250a and Mtf1 (See quantification in Fig. 4F). Each panel presents an overview of immunostained cells (upper panel, scale bar = 10 μm) and a zoomed-in view of a highlighted cell (lower panel, scale bar = 5 μm).

#### SUPPLEMENTAL FIGURE 7

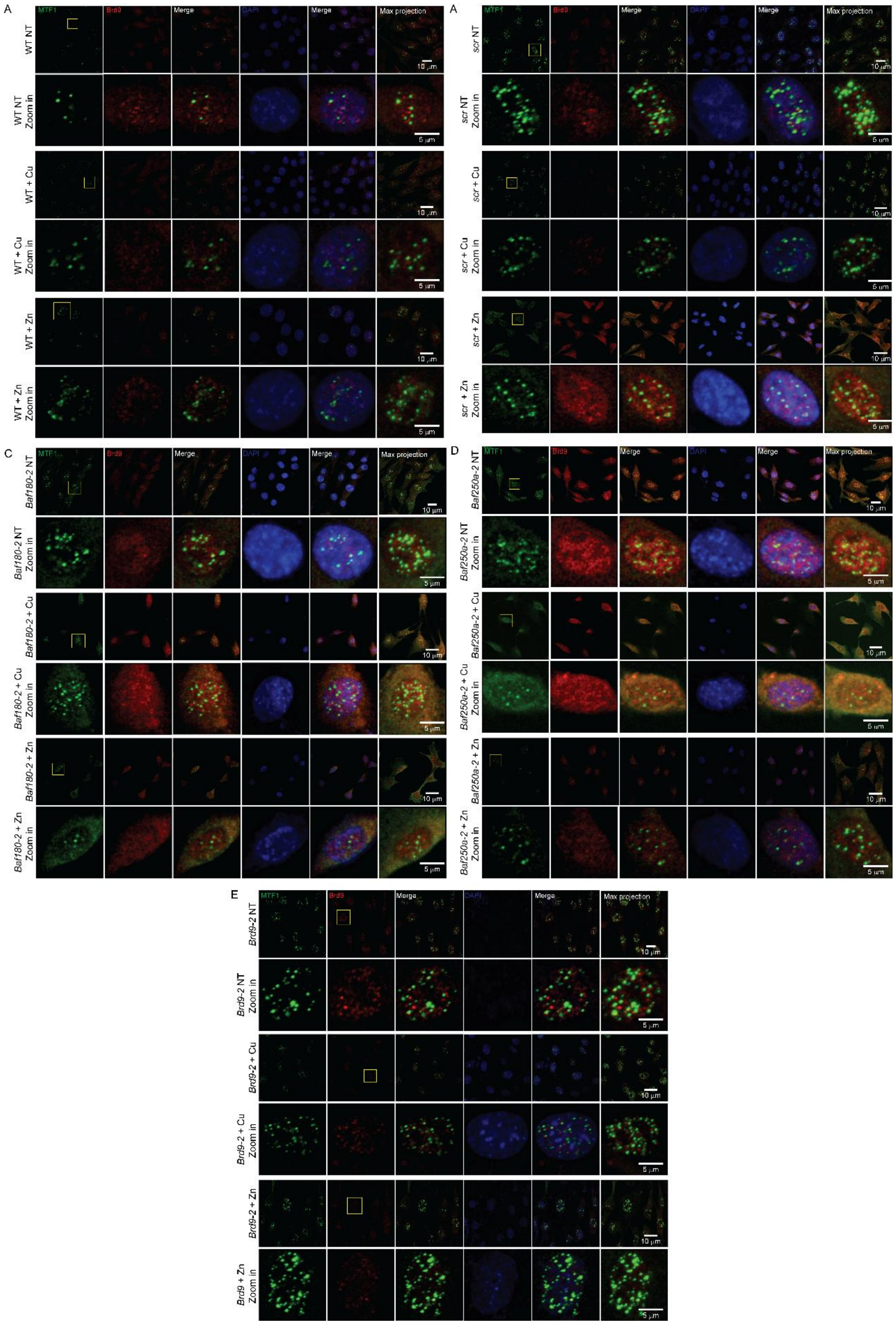

**Supplemental Figure 7: Expression and nuclear localization of Brd9 in proliferating C2C12 myoblasts.** Representative confocal microscopy images showing the expression and localization of Brd9 (red), Mtf1 (green), and DAPI (blue) in proliferating wild type (A), scr (B), *Baf180* KD (C), *Baf250a* KD (D) and *Brd9* KD (E) C2C12 myoblasts. Nuclei are counterstained with DAPI (blue). Non-treated myoblasts exhibit nuclear Brd9 localization, with no colocalization observed between Brd9 and Mtf1 (See quantification in Fig. 4F). Each panel presents an overview of immunostained cells (upper panel, scale bar = 10 μm) and a zoomed-in view of a highlighted cell (lower panel, scale bar = 5 μm).

#### SUPPLEMENTAL FIGURE 8

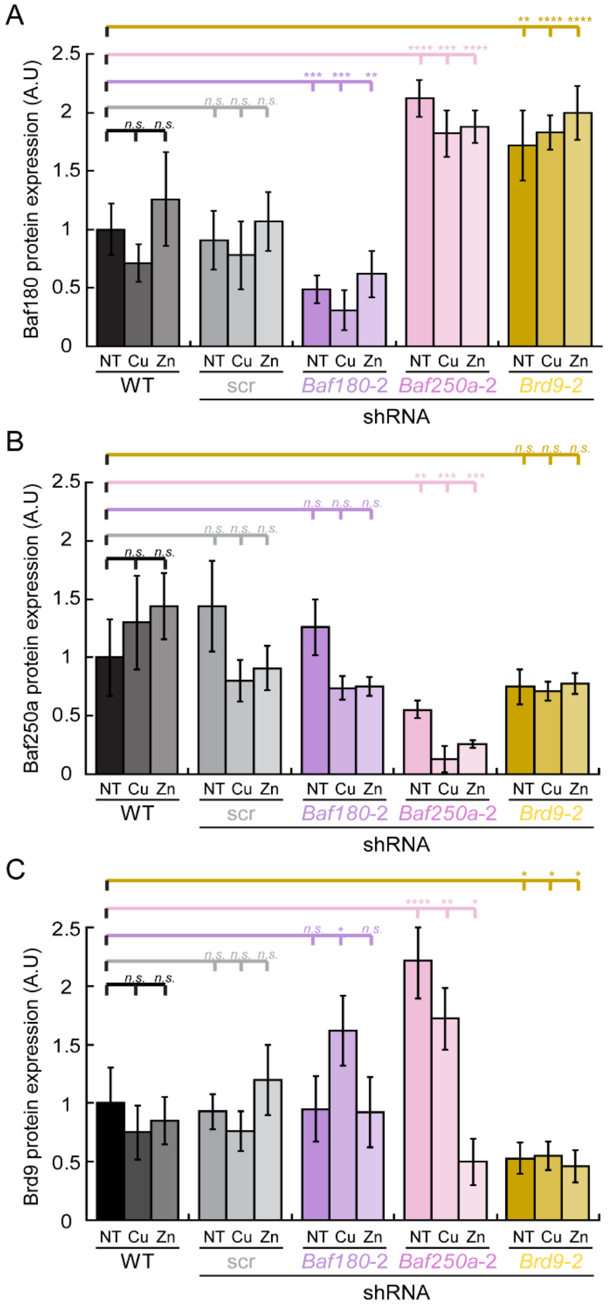


**Supplemental Figure 8. Quantification of SWI/SNF subunit protein levels in proliferating C2C12 myoblasts.** Confocal microscopy-based quantification of protein expression for **(A)** Baf180, **(B)** Baf250a, and **(C)** Brd9 in wild type (WT), scr, *Baf180* KD, *Baf250a* KD, and *Brd9* KD myoblasts under different conditions: Untreated (NT), $CuSO_4$ (100 μM, 48h) and $ZnSO_4$ treatment (50 μM, 48h). Baf180 levels are increased in *Baf250a* KD and *Brd9* KD myoblasts, suggesting potential compensatory regulation. Data represents the mean ± SE of three independent biological replicates. *P < 0.05, **P < 0.01, ****P < 0.0001.

**SUPPLEMENTAL FIGURE 9**

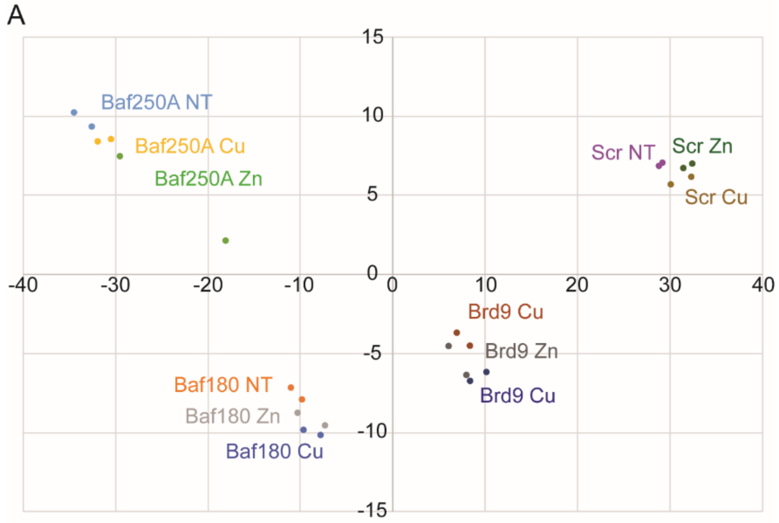


**Supplemental Figure 9. Principal component analysis reveals consistent global transcriptome profiles across conditions.** Principal Component Analysis (PCA) comparing global gene expression profiles of proliferating Scr control, and *Baf180*, *Baf250a*, and *Brd9* KD myoblasts, under both untreated conditions and following 48-hour treatment with 100 µM $CuSO_4$ or 50 µM $ZnSO_4$. PCA clustering indicates that transcriptomic profiles remain consistent within each knockdown condition, regardless of metal treatment.

#### SUPPLEMENTAL FIGURE 10

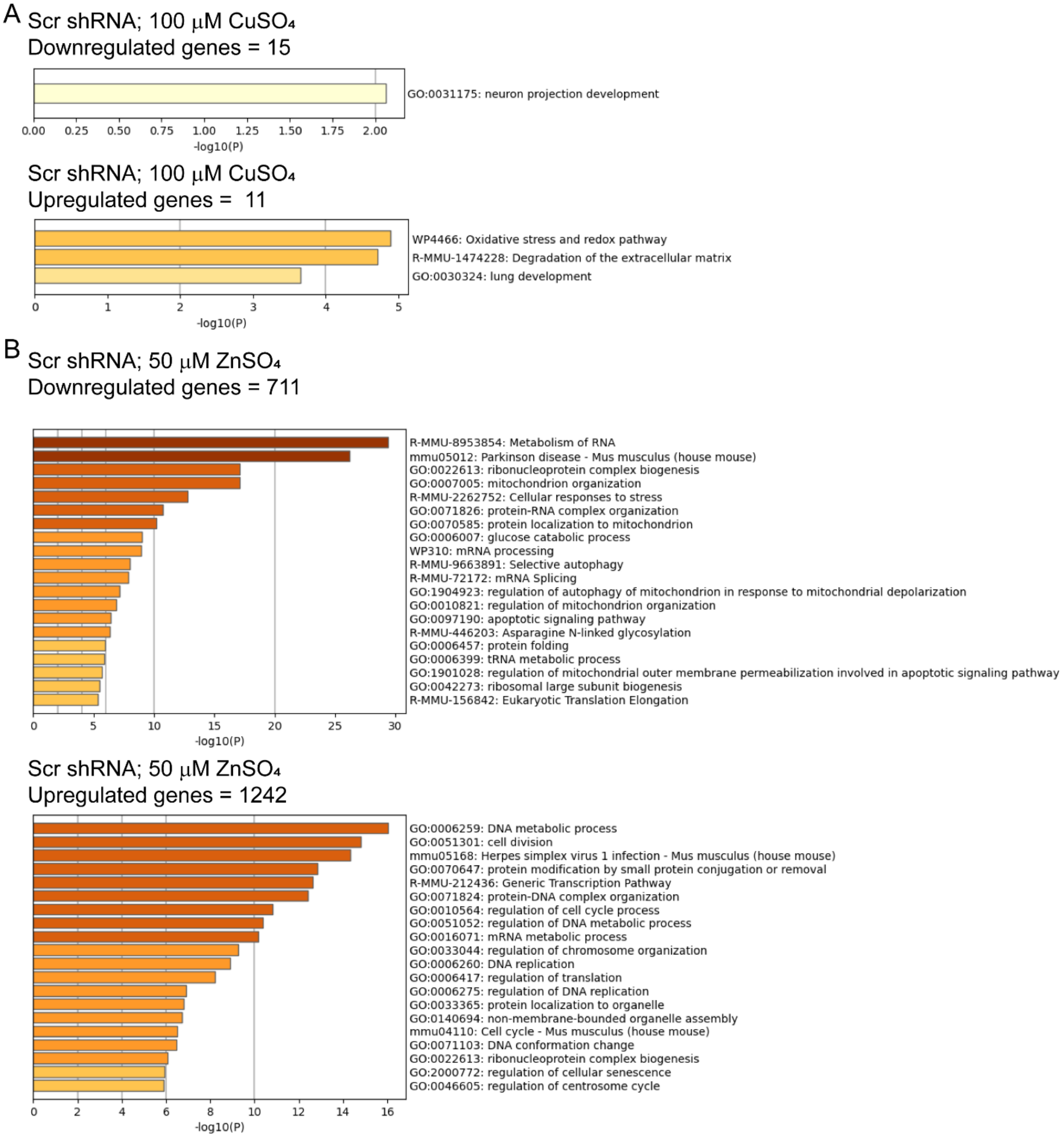


**Supplemental Figure 10. GO analysis of DEGs in scr control myoblasts supplemented with metals.** DEG identified from scr control cells cultured in proliferation media supplemented with 100 μM $CuSO_4$ **(A)** or 50 μM $ZnSO_4$ (B) were compared to the same cell line cultured in the absence of metals (basal media). The patterns represent downregulation and upregulation of DEGs shown in **Supp. Table 4**.

**SUPPLEMENTAL FIGURE 11**

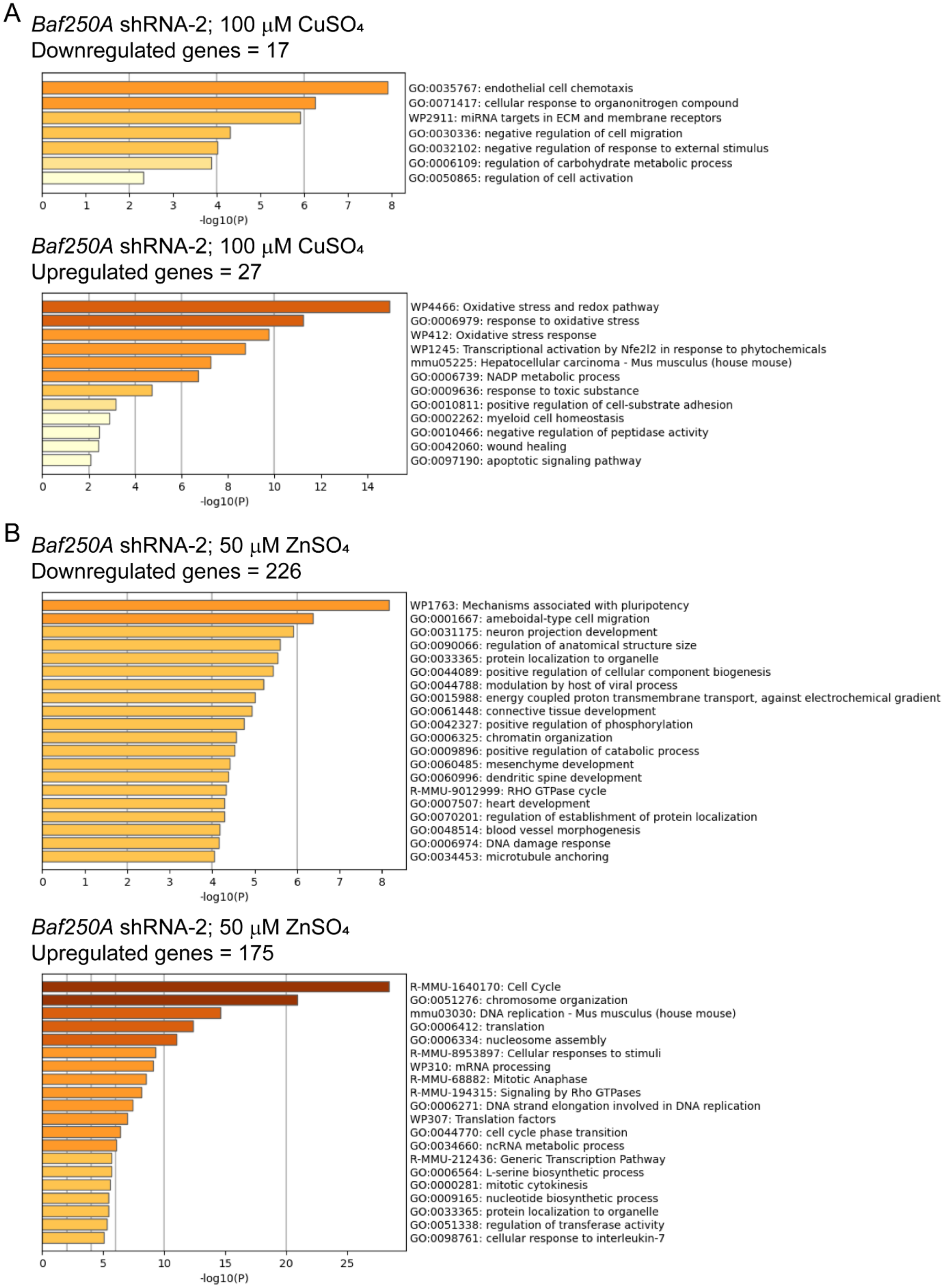


**Supplemental Figure 11.** GO analysis of DEGs in *Baf250a* KD myoblasts supplemented with metals. DEG identified from *Baf250a* KD cells cultured in proliferation media supplemented with 100 µM $CuSO_4$ **(A)** or 50 µM $ZnSO_4$ **(B)** were compared to the same cell line cultured in the absence of metals (basal media). The patterns represent downregulation and upregulation of DEGs shown in **Supp. Table 4**.

#### SUPPLEMENTAL FIGURE 12

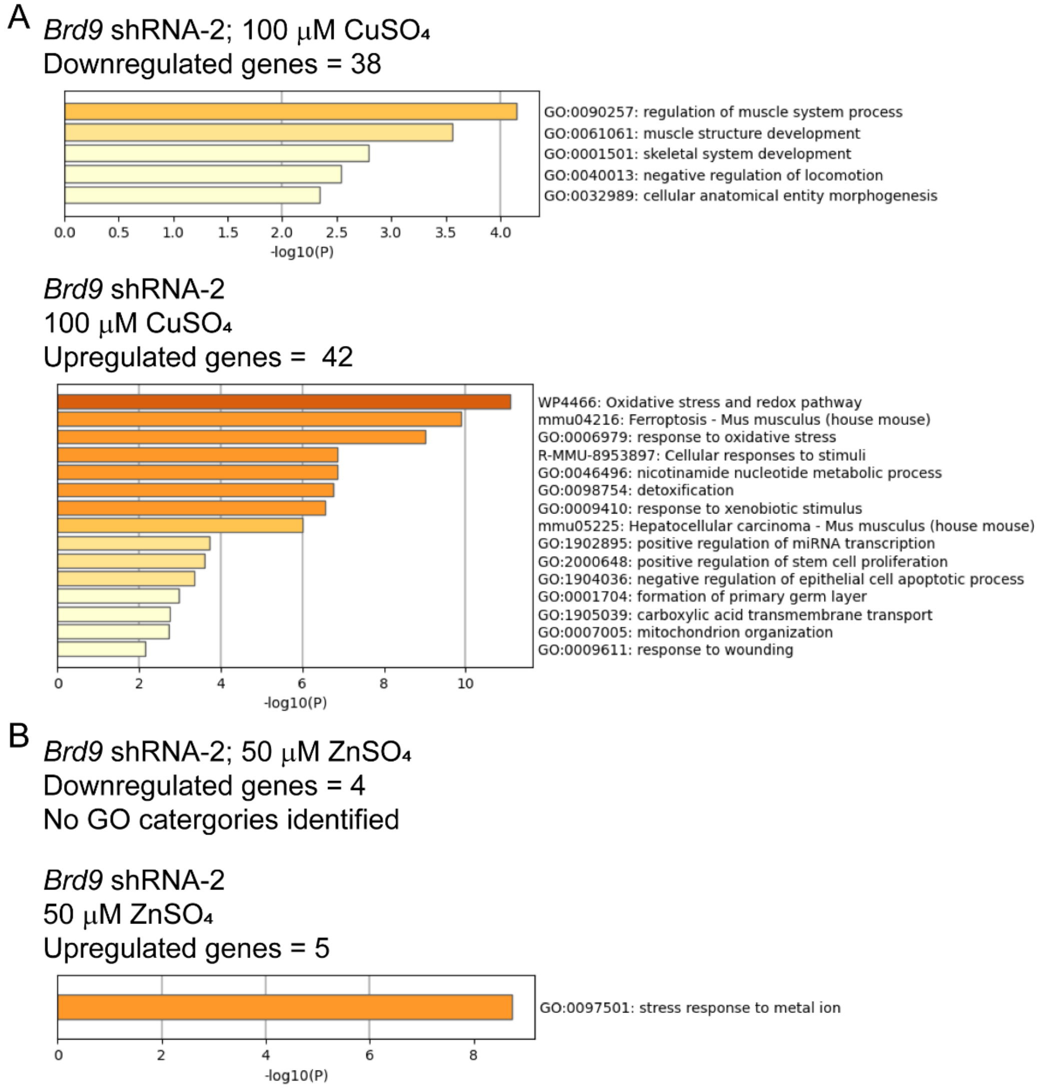


**Supplemental Figure 12.** GO analysis of DEGs in *Brd9* KD myoblasts supplemented with metals. DEG identified from *Brd9* KD cells cultured in proliferation media supplemented with 100 μM $CuSO_4$ **(A)** or 50 μM $ZnSO_4$ (B) were compared to the same cell line cultured in the absence of metals (basal media). The patterns represent downregulation and upregulation of DEGs shown in **Supp. Table 4**.

### SUPPLEMENTAL FIGURE 13

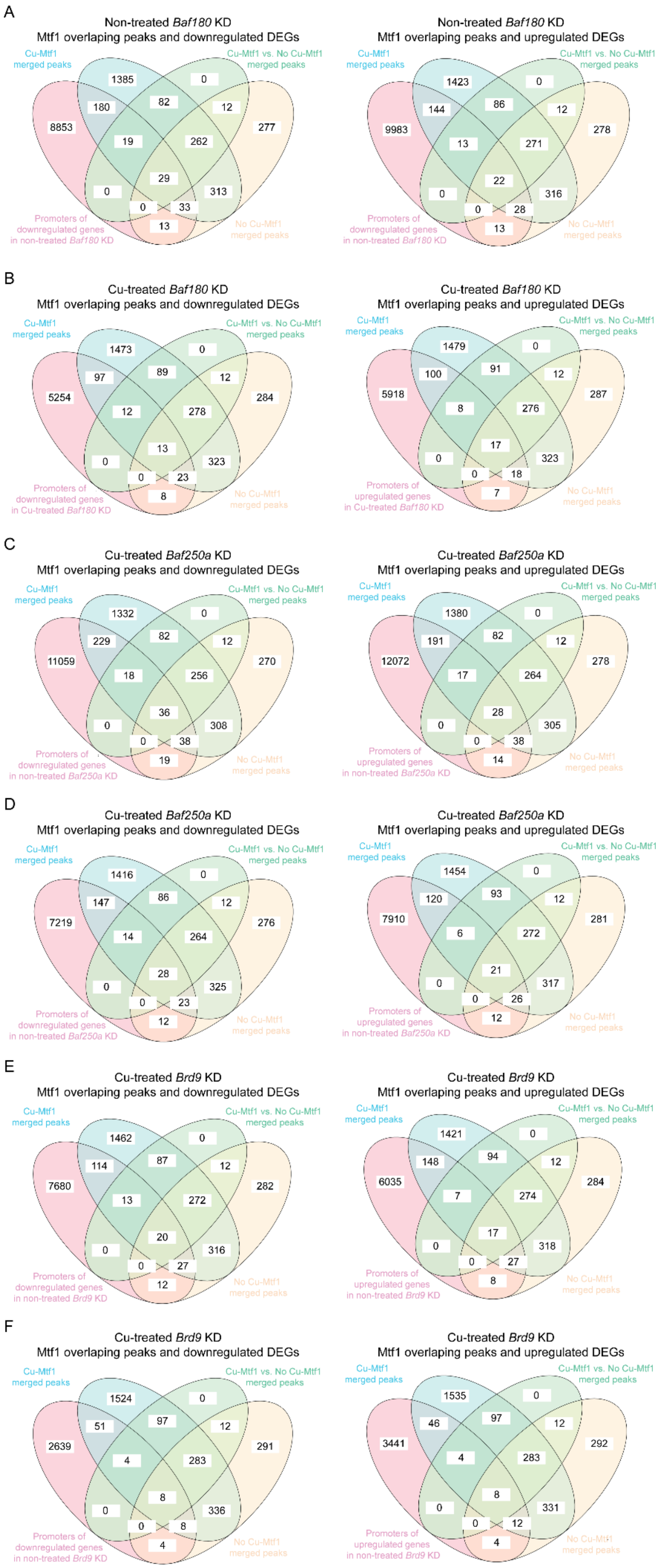

**Supplemental Figure 13. MTF1 chromatin binding correlates with DEGs in SWI/SNF KD myoblasts under Cu exposure.** The Venn diagrams illustrate the overlap of differentially expressed genes (DEGs) that are either upregulated or downregulated across *Baf180* **(A, B)**, *Baf250a* **(C, D)**, and *Brd9* **(E, F)** KD myoblasts, cultured with or without Cu treatment. These transcriptional changes were further integrated with MTF1 chromatin occupancy data obtained from CUT&RUN analysis of non-treated and Cu-treated cells, providing insight into the regulatory landscape influenced by MTF1. Notably, a subset of DEGs across the different SWI/SNF subunit KD conditions overlapped with MTF1 binding sites, suggesting that MTF1 directly regulates a fraction of genes affected by SWI/SNF disruption. The shared and unique gene sets highlight the interplay between SWI/SNF chromatin remodelers and MTF1 in orchestrating transcriptional responses to Cu, reinforcing the role of these factors in metal-responsive gene regulation and myoblast adaptation to metal stress.

### SUPPLEMENTAL TABLES

**Supplemental Table 1. Sequences of shRNA used in this study (2, 3).**

| | | |
|---|---|---|
| *Baf250A* shRNA1 | CCGGCTTTATAGTATGGCGAGTTAACTCGAGTTAACTCGCCAT ACTATAAAGTTTTTG | TRCN0000238304 |
| *Baf250A* shRNA2 | CCGGCCTAGGCAGCCTAACTATAATCTCGAGATTATAGTTAGG CTGCCTAGGTTTTTG | TRCN0000238306 |
| *Brd9* shRNA1 | CCGGTGGACTTTGGCACGATGAAAGCTCGAGCTTTCATCGT GCCAAAGTCCATTTTTG | TRCN0000225737 |
| *Brd9* shRNA2 | CCGGCACCGAATGGTGTCCAATAAGCTCGAGCTTATTGGACA CCATTCGGTGTTTTTG | TRCN0000225739 |
| *Baf180* shRNA1 | CCGGTGTGAAGTTGGTCCTAGTTTACTCGAGTAAACTAGGAC CAACTTCACATTTTTG | TRCN0000304680 |
| *Baf180* shRNA2 | CCGGGTGCAATATCCAGACTATTATCTCGAGATAATAGTCTGG ATATTGCACTTTTTG | TRCN0000304681 |
| scr shRNA | CCGGCAACAAGATGAAGAGCACCAACTCGAGTTGGTGCTCT TCATCTTGTTGTTTTT | pLKO.1-puro non-target shRNA control plasmid DNA MFCD07785395 SHC002 |

**Supplemental Table 4. Summary of DEG genes from metal-treated KD and scr control cells.** Data shows the differences within each strain upon metal supplementation

| Baf180 KD | | | Baf250a KD | | | Brd9 KD | | | SCR | | |
|---|---|---|---|---|---|---|---|---|---|---|---|
| +Cu vs NT | DEGs | % DE genes | +Cu vs NT | DEGs | % DE genes | +Cu vs NT | DEGs | % DE genes | +Cu vs NT | DEGs | % DE genes |
| Sig. up | 1308 | 0.044 | Sig. up | 28 | 0.001 | Sig. up | 42 | 0.0016 | Sig. up | 11 | 0.00037 |
| Sig. down | 1510 | 0.051 | Sig. down | 17 | 0.00064 | Sig. down | 39 | 0.0014 | Sig. down | 19 | 0.00064 |
| All sig. | 2818 | 0.095 | All sig. | 45 | 0.00164 | All sig. | 81 | 0.003 | All sig. | 30 | 0.00101 |
| +Zn vs NT | DEGs | % DE genes | +Zn vs NT | DEGs | % DE genes | +Zn vs NT | DEGs | % DE genes | +Zn vs NT | DEGs | % DE genes |
| Sig. up in KO | 137 | 0.0046 | Sig. up in KO | 176 | 0.0066 | Sig. up in KO | 5 | 0.00019 | Sig. up in KO | 1261 | 0.043 |
| Sig. down in KO | 324 | 0.011 | Sig. down in KO | 227 | 0.0085 | Sig. down in KO | 4 | 0.00015 | Sig. down in KO | 718 | 0.024 |
| All sig. | 461 | 0.0156 | All sig. | 403 | 0.0151 | All sig. | 9 | 0.00034 | All sig. | 1979 | 0.067 |
